# NLRP3, NLRP6, and NLRP12 are inflammasomes with distinct expression patterns

**DOI:** 10.1101/2024.02.05.579000

**Authors:** Bo Wei, Zachary P. Billman, Kengo Nozaki, Helen S. Goodridge, Edward A. Miao

## Abstract

Inflammasomes are sensors that detect cytosolic microbial molecules or cellular damage, and in response they initiate a form of lytic regulated cell death called pyroptosis. Inflammasomes signal via homotypic protein-protein interactions where CARD or PYD domains are crucial for recruiting downstream partners. Here, we screened these domains from NLR family proteins, and found that the PYD domain of NLRP6 and NLRP12 could activate caspase-1 to induce cleavage of IL-1β and GSDMD. Inflammasome reconstitution verified that full length NLRP6 and NLRP12 formed inflammasomes in vitro, and NLRP6 was more prone to auto-activation. NLRP6 was highly expressed in intestinal epithelial cells (IEC), but not in immune cells. Molecular phylogeny analysis found that NLRP12 was closely related to NLRP3, but the activation mechanisms are different. NLRP3 was highly expressed in monocytes and macrophages, and was modestly but appreciably expressed in neutrophils. In contrast, NLRP12 was specifically expressed in neutrophils and eosinophils, but was not detectable in macrophages. NLRP12 mutations cause a periodic fever syndrome called NLRP12 autoinflammatory disease. We found that several of these patient mutations caused spontaneous activation of caspase-1 in vitro, which likely causes their autoinflammatory disease. Different cell types have unique cellular physiology and structures which could be perturbed by a pathogen, necessitating expression of distinct inflammasome sensors to monitor for signs of infection.

## INTRODUCTION

Pyroptosis is a form of regulated cell death that is proinflammatory and uniquely gasdermin-dependent (*1*). Pyroptosis exhibits large membrane balloons and occurs concomitantly with release of the proinflammatory cytokines IL-1β and IL-18. Inflammasomes are the sensors that most commonly initiate pyroptosis (*2*). Pathogen-associated molecular patterns (PAMPs) and host cell generated danger-associated molecular patterns (DAMPs) are recognized by germline encoded pattern recognition receptors (PRRs) (*3*). Then the PRRs oligomerize and activate the downstream protease caspase-1 directly or through the adapter protein ASC (*2*). Multiprotein oligomerized platforms that activate caspase-1 are called inflammasomes. Once caspase-1 is activated, it cleaves the inflammatory cytokines pro-IL-1β and pro-IL-18 to their mature forms and additionally cleaves gasdermin D (GSDMD) (*1, 2*). After cleavage, the N-terminus of GSDMD moves to the plasma membrane, oligomerizes, and forms pores in the membrane with an inner diameter of 10 to 14 nm (*1, 4, 5*). These pores eventually cause pyroptosis. In parallel, caspase-4/5/11 act as sensors for cytosolic LPS and also cause pyroptosis (*6–8*). Pyroptosis is quite effective in preventing infection by environmental pathogens, and many host-adapted pathogens use virulence factors to inhibit pyroptosis in order to facilitate intracellular replication.

Bacterial infection can be sensed when type 3 secretion systems (T3SS) translocate flagellin, rod, or needle protein into the host cell cytosol. These three agonists directly bind to NAIP sensors that induce the oligomerization of NLRC4. This NLRC4 oligomer can directly interact with and activate caspase-1; additionally, this signaling can also be amplified through the adaptor ASC (*9*). In contrast, the NLRP3 inflammasome assembles in response to various stimuli including ATP, nigericin, and crystals (*10*). These stimuli cause cellular perturbations that, through mechanisms still being described, cause NLRP3 to oligomerize. This oligomerization requires help from NEK7, which acts as a structural co-factor that binds to NLRP3 (*11–13*). Unlike the NLRC4 inflammasome, NLRP3 oligomers cannot activate caspase-1 directly, but must signal through the adapter protein ASC.

Both NLRC4 and NLRP3 belong to the NLR superfamily. Based on differences in their N-terminus, NLR proteins are divided into NLRA, NLRB, NLRC, NLRP, and NLRX subfamilies (*14*). Of these, several NLRs form inflammasomes, including NLRB, NLRC, and NLRP family members, which contain N-terminal BIR domains, caspase recruitment domains (CARDs), or pyrin domains (PYDs), respectively. NAIPs are in the NLRB family, however the BIR domains do not bind to downstream signaling proteins, instead the co-polymerizing NLRC4 contains the CARD domain of the NAIP/NLRC4 inflammasome (*9*). NLRP3 contains a PYD domain that confers downstream signaling to this inflammasome (*2*).

Interactions between NLRs, the adaptor protein ASC, and caspase-1 are mediated by homotypic protein-protein interactions of their CARD or PYD domains, which are members of the death domain superfamily (*15*). Thus, a specific CARD will interact with another cognate CARD (CARD-CARD interactions); similarly, homotypic PYD-PYD interactions also occur (*15*). However, not every CARD interacts with every other CARD, rather, the interactions are specific. For example, the CARD of NLRC4 interacts with the CARD of ASC or the CARD of caspase-1 (*9*), but not with the CARD of the type I interferon signaling protein MAVS. As an adaptor protein, ASC contains both a PYD and a CARD that bridges upstream PYD-containing sensors to the CARD of caspase-1. Additionally, ASC can also be recruited to CARD-containing sensors to amplify their signaling. While the protein-protein interactions of the NLRC4 and NLRP3 inflammasomes have been well-characterized, the interactions for other NLR family members remain poorly defined. Here, we search for additional inflammasomes by screening NLR family members for their ability to signal to ASC and/or caspase-1.

## RESULTS

### Identify CARD and PYD domains that activate caspase-1

We wondered whether other proteins that also contain CARD or PYD could activate caspase-1 via homotypic protein-protein interactions. We chose to use the chemically inducible dimerization system, where protein-protein interactions are specifically induced by addition of dimerizing drug (Fig. 1A). The CARD and PYD domains of interest were fused to the DmrB binding domain, which originates from the FK506-binding protein FKBP12. Homo-dimerization is induced by the cell-permeable ligand AP20187 (also called B/B homodimerizer). This method has been previously used to study inflammasome signaling (*16*), where DmrB dimerization simulates the oligomerization of an NLR inflammasome. We first optimized our transfection conditions to avoid autoactivation of ASC and caspase-1.

**Fig. 1.**
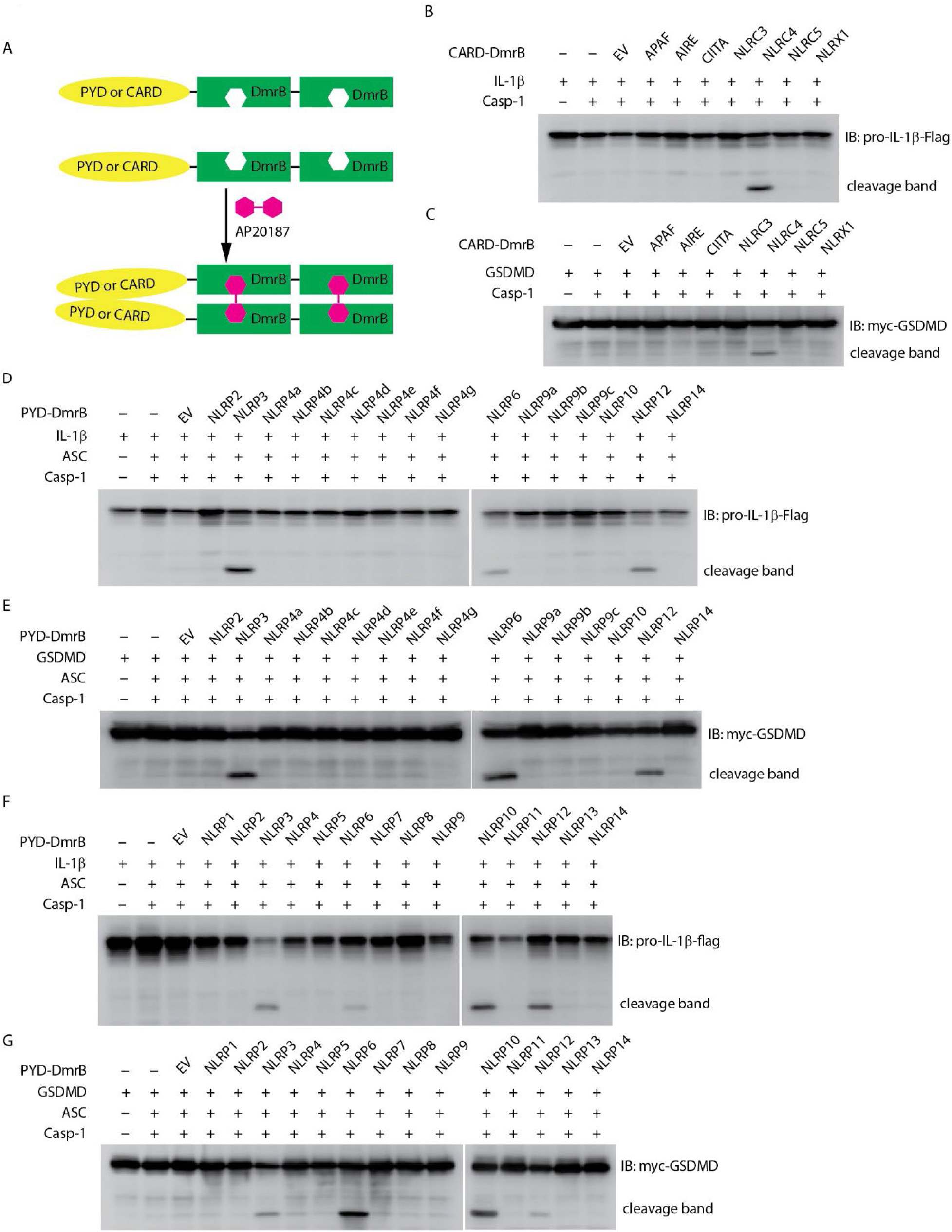
Screening CARD and PYD domains by inducible dimerization. (A) Diagram of AP20187 induced dimerization experiment system. (B-G) HEK293T/17 cells were transiently transfected with indicated plasmids. 24 hours later the dimer inducer AP20187 (20 nM) was added for another 10 hours. Cells were harvested and lysed, then lysates were subjected to immunoblotting with indicated antibody. EV, empty vector. (B and C) Mouse CARD domain fusion proteins did not induce cleavage of IL-1β (B) and GSDMD (C) through Caspase-1. The CARD domain from *Apaf*, *Aire*, *Ciita*, *Nlrc3*, and *Nlrc4*, as well as the N-terminal domain of *Nlrx1* were expressed as fusion proteins with tandem DmrB domains. (D and E) Mouse NLRP6 and NLRP12 PYD domain fusion proteins induced cleavage of IL-1β (D) and GSDMD (E). The PYD domain from *Nlrp2*, *Nlrp3*, *Nlrp4a, Nlrp4b, Nlrp4c, Nlrp4d, Nlrp4e*, *Nlrp4f*, *Nlrp4g*, *Nlrp6*, *Nlrp9a*, *Nlrp9b, Nlrp9c, Nlrp10*, *Nlrp12*, and *Nlrp14* were expressed as fusion proteins with tandem DmrB domains. (F and G) Human NLRP6, NLRP10 and NLRP12 PYD domain fusion proteins induced cleavage of IL-1β (F) and GSDMD (G). The PYD domain from *NLRP1*, *NLRP2*, *NLRP3*, *NLRP4*, *NLRP5*, *NLRP6*, *NLRP7*, *NLRP8*, *NLRP9*, *NLRP10*, *NLRP11*, *NLRP12*, *NLRP13*, and *NLRP14* were expressed as fusion proteins with tandem DmrB domains. All the blotting results are representative of at least 3 independent experiments.

We first examined proteins with CARD domains to determine whether they activate caspase-1. We transfected HEK293T/17 cells with IL-1β, caspase-1, and the CARD-DmrB construct, and assessed IL-1β cleavage by western blotting. We used the CARD domain of NLRC4 as a positive control (*9*), and confirmed that fusion of the NLRC4 CARD to DmrB and dimerization with AP20187 activates caspase-1 and causes IL-1β cleavage (Fig. 1B). In the absence of AP20187 we observed notably weaker IL-1β cleavage (data not shown). We next tested the CARD domains from other proteins that are not known to activate caspase-1: APAF, AIRE, CIITA, NLRC3, and NLRC5 (*14*) (Fig. S1A). In contrast to NLRC4, none of these CARD-DmrB fusion proteins induced cleavage of IL-1β (Fig. 1B), though they were all expressed at relatively similar levels (Fig. S1A). Although it is not classified as a CARD domain, we also tested the N-terminal domain of NLRX1, which also failed to induce IL-1β cleavage (Fig. 1B). Similarly, the processing of GSDMD was only induced by the NLRC4 CARD fusion protein, but none of other proteins (Fig. 1C).

Next, we examined PYD domains from mouse NLRP proteins (*14*) (Fig. S1B). In the classical NLRP3 inflammasome, NLRP3 recruits the adaptor protein ASC through PYD-PYD interactions (*2*). We used the NLRP3 PYD as a positive control and verified that dimerization accomplishes IL-1β cleavage (Fig. 1D). We next tested the PYD domains of all other NLRP proteins from mice (Fig. S1B). Interestingly, the NLRP6 PYD and the NLRP12 PYD fusion proteins also induced IL-1β cleavage, whereas the other PYD fusion proteins which originated from NLRP2, NLRP4a, NLRP4b, NLRP4c, NLRP4d, NLRP4e, NLRP4f, NLRP4g, NLRP6, NLRP9a, NLRP9b, NLRP9c, NLRP10, or NLRP14 did not cause any cleavage of IL-1β (Fig. 1D). Similarly, only the NLRP3, NLRP6, and NLRP12 PYD fusion proteins resulted in cleavage of GSDMD (Fig. 1E). To our surprise, our results did not validate two recently described inflammasomes. NLRP9b has been reported to recognize short double-stranded RNA stretches via RNA helicase DHX9 and form inflammasome complexes together with the adaptor proteins ASC and caspase-1 (*17*). NLRP10 has been reported to monitor mitochondrial integrity in an mtDNA-independent manner, and form inflammasome with ASC to activate caspase-1 (*18, 19*). However, their PYD domains did not result in IL-1β cleavage or GSDMD cleavage in our inducible dimerization system. Therefore, this system may be susceptible to false-negative results.

We next studied the PYD domains from human NLRP proteins (*14*) (Fig. S1C). Similar to the results from the mouse homologs, the PYD from NLRP3, NLRP6, and NLRP12 resulted in cleavage of IL-1β and GSDMD, while neither IL-1β nor GSDMD were inducibly cleaved by NLRP1, NLRP2, NLRP4, NLRP5, NLRP7, NLRP8, NLRP9, NLRP11, NLRP13, or NLRP14 (Fig. 1F-1G). To our surprise again, the human NLRP7 PYD did not result in cleavage of IL-1β, although NLRP7 has been reported to recognize microbial lipopeptides in human macrophage and assemble inflammasome (*20*). Interestingly, we found human NLRP10 PYD fusions did result in cleavage of IL-1β and GSDMD, whereas the PYD domain of mouse NLRP10 could not (Fig. 1D-1G). This difference may be due to false-negative results with the murine NLRP10, or could be caused by the absence of murine co-factors that are essential but which do not exist in human HEK293T/17 cells. PYD and CARD domains are highly diverse with no specific identifying sequence or motif, and thus individual PYD or CARD domains could be subject to specific regulatory modification. For example, the NLRP3 PYD domain can be post translationally modified by phosphorylation, acetylation, or other modifications (*21, 22*). It may be that an absent posttranslational modification of PYD domain of NLRP7, NLRP9b, and mouse NLRP10 caused a false negative result in our assay.

Taken together, we observed that in addition to NLRP3, the PYD domains of NLRP6, NLRP10, and NLRP12 activate caspase-1 and result in cleavage of IL-1β and GSDMD. We chose to further study NLRP3, NLRP6, and NLRP12.

### Reconstitution of the NLRP3 inflammasome

NLRP3 inflammasome signaling can be reconstituted in HEK293T cells (*23*), and we wanted to use this approach to study NLRP6, and NLRP12. First, we optimized the NLRP3 reconstitution as a positive control. We first choose to detect cytosolic IL-β cleavage in live cells by western blot. We optimized the system with NLRP3, and used optimized transfection conditions where ASC and caspase-1 alone did not cause autoactivation (Fig. 2A). Expression of full length of NLRP3 together with ASC and caspase-1 resulted in cleavage of IL-1β by western blot (Fig. 2A). The amount of NLRP3 we transfected resulted in autoactivation without application of specific NLRP3 agonists (Fig. 2A).

**Fig. 2.**
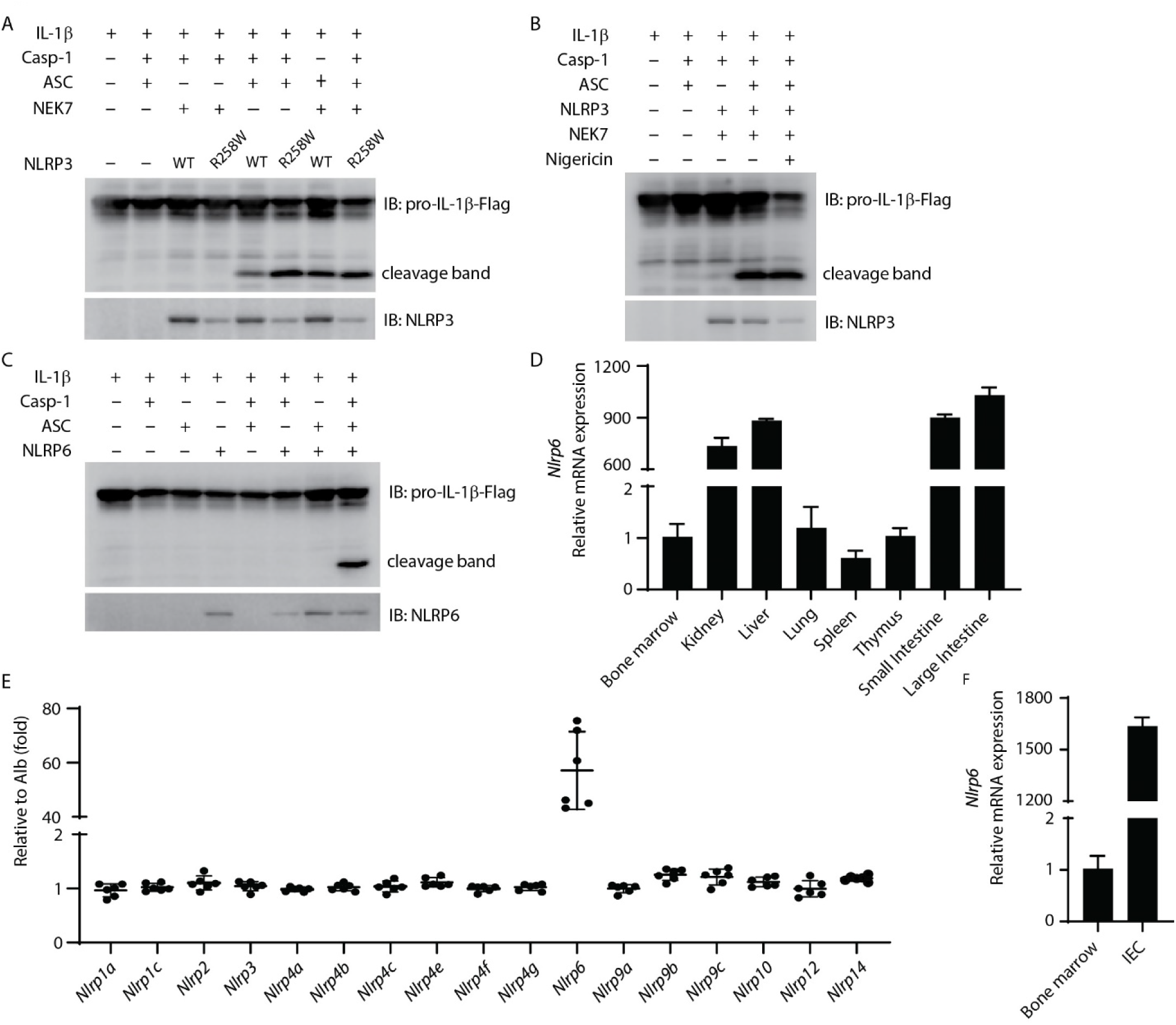
Reconstitution of mouse NLRP3 and NLRP6 inflammasome in vitro. (A-B) Reconstitution of the NLRP3 inflammasome in HEK293T/17 cells with IL-1β, caspase-1, ASC, NEK7, and either NLRP3 (WT) or NLRP3(R258W). (B) Addition of nigericin (20 μM) treatment for 90min. Cleaved IL-1β was detected by western blot. (C) Reconstitution of the NLRP6 inflammasome in HEK293T/17 cells with IL-1β, caspase-1, ASC, and NLRP6. Cleaved IL-1β was detected by western blot. (D) qRT-PCR analysis of *Nlrp6* expression in indicated mouse tissues. (E) The relative expression of *Nlrp* genes in IECs. Data extracted from the original data in NCBI (GDS3921). The expression results were normalized by albumin (*Alb*) expression as a gene that should not be expressed (thus a value of 1 reflects absent expression). (F) qRT-PCR analysis of *Nlrp6* expression in purified IECs compared to bone marrow cells. All the results are representative of at least 3 independent experiments.

We next validated that the NLRP3 hyperactivation mutant R258W resulted in enhanced cleavage of IL-1β compared to WT NLRP3 (*24*) (Fig. 2A). Interestingly, ectopic expression of NEK7 increased the cleavage of IL-1β in cells expressing WT NLRP3 (Fig. 2A, lane 5 and lane 7), indicating that endogenous NEK7 in HEK293T/17 cells was sufficient, but could be enhanced by overexpression. Such overexpression of NEK7 did not enhance IL-1β cleavage in cells expressing NLRP3(R258W) (Fig. 2A, lane 6 and lane 8).

Reconstituted NLRP3 in HEK293T/17 cells stimulated with nigericin has been published to cause IL-1β release detectable by ELISA (*23*). When we assayed IL-1β release by ELISA, we did observe a nigericin-dependent IL-1β ELISA signal. However, we also observed ASC independent IL-1β release (Fig. S2A), and notably HEK293T/17 cells do not express GSDMD that normally releases IL-1β from the cell. Addition of the NLRP3 agonist nigericin did not further increase cleavage of IL-1β in these cells by western blot (Fig. 2B). These results suggest that this ELISA signal was a consequence of nigericin toxicity rather than NLRP3 signaling. Therefore, we did not continue to use nigericin or ELISA in our reconstitution experiments in HEK293T/17 cells.

### NLPR6 is expressed in intestinal epithelial cells and is prone to autoactivation

We used the same HEK293T/17expression system to study NLRP6, which was transfected together with ASC, caspase-1, and IL-1β. Remarkably, when NLRP6 was cotransfected with ASC and caspase-1, it resulted in pronounced cleavage of IL-1β (Fig. 2C). Although caspase-11 has been published to promote NLRP6 signaling (*25*), when we added caspase-11 to the transfection, this did not enhance IL-1β cleavage (Fig. S2B). Similarly, NEK7 overexpression did not enhance NLRP6 signaling (Fig. S2C). Therefore, NLRP6 overexpression results in its autoactivation.

We next examined *Nlrp6* expression in several gene expression databases (BioGPS, ImmGen, and the Mouse Cell Atlas). *Nlrp6* appears to not be expressed in immune cells (Fig. S2D, S2E, and S2F). qPCR results from cell lines also showed that *Nlrp6* was poorly expressed in macrophages (BMDMs, RAW264.7, J774A.1), and *NLRP6* was poorly expressed in human immune cell lines (HL-60, THP1, U937) and human epithelial cell lines (Caco2, T84) (Table S1).

In contrast, results from tissue samples showed high *Nlrp6* expression in the intestine, but not in the spleen, bone marrow, or blood (Fig. 2D). In support of this, a publicly available dataset from Reikvam *et al.* analyzing purified intestinal epithelial cells (IECs) showed that among mouse *Nlrp* genes, only *Nlrp6* was highly expressed in IECs (*26*) (Fig. 2E). To confirm expression in IECs, we purified IECs from the intestine and isolated RNA in comparison to RNA from bone marrow. *Nlrp6* was indeed expressed strongly in the IECs with negligible signal from bone marrow cells (Fig. 2F). Interestingly, in the Reikvam *et al.* dataset *Nlrc4* and *Aim2* were also appreciably expressed (Fig. S2G), but at lower levels than *Nlrp6*. Overall, the NLRP6 inflammasome is expressed in IECs, but appears to not be expressed in immune cells.

### NLRP12 is an inflammasome that is less prone to autoactivation

To study the function of NLRP12, we set up a similar in vitro activation assay in HEK293T/17 cells as used above. To our surprise, expression of full length NLRP12 did not induce IL-1β cleavage via ASC and caspase-1 (Fig. 3A), suggesting that the protein is less prone to autoactivation compared to NLRP3 and NLRP6. The addition of NEK7 did not cause autoactivation of NLRP12 (Fig. S3A).

**Fig. 3.**
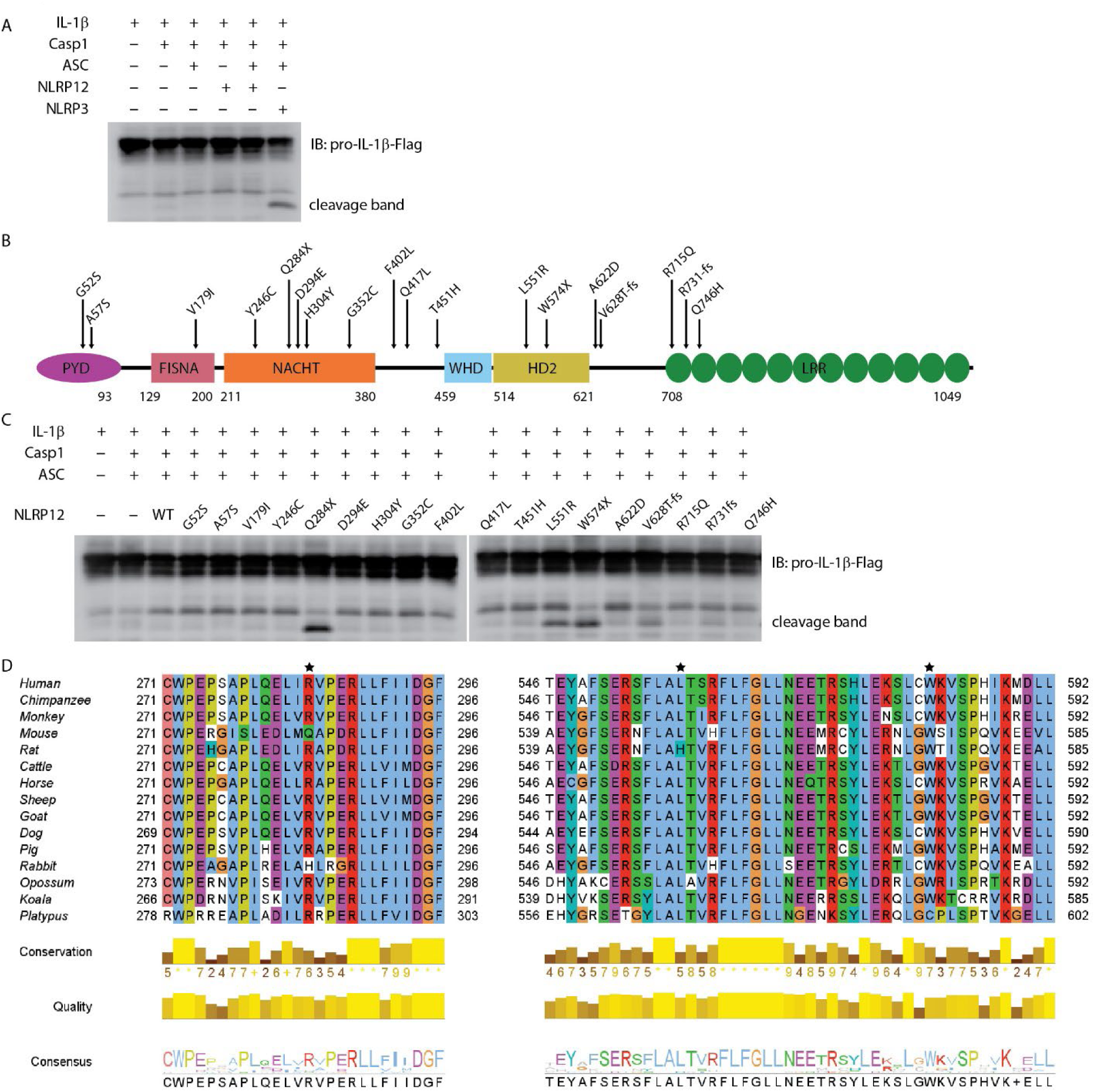
Reconstitution of NLRP12 inflammasome in vitro. (A) Reconstitution of NLRP12 inflammasome in HEK293T/17 cells with IL-1β, Caspase-1, ASC, NLRP12 (WT), and NLRP3 (as a positive control). Cleaved IL-1β was detected by western blot. (B) Schematic diagram of mouse NLRP12 mutants based on human NLRP12 mutants reported in human NLRP12-AID patients. (C) Reconstitution of NLRP12 inflammasome with IL-1β, caspase-1, ASC, NLRP12 (WT), and various NLRP12 mutants. Cleaved IL-1β was detected by western blot. (D) Alignment of NLRP12 protein sequence from indicated species. Asterisks indicate the mouse amino acids Gln^284^, Leu^551^ and Trp^574^ that correspond to residues mutated in NLRP12-AID patients (human Arg^284^, Leu^558^ and Trp^581^). Human, *Homo sapiens*; Chimpanzee, *Pan troglodytes*; Monkey, *Macaca mulatta*; Mouse, *Mus musculus*; Rat, *Rattus norvegicus*; Cattle, *Bos taurus*; Horse, *Equus caballus*; Sheep, *Ovis aries*; Goat, *Capra hircus*; Dog, *Canis lupus familiaris*; Pig, *Sus scrofa*; Rabbit, *Oryctolagus cuniculus*; Opossum, *Monodelphis domestica*; Koala, *Phascolarctos cinereus*; Platypus, *Ornithorhynchus anatinus*. All the results are representative of at least 3 independent experiments.

Some patients with periodic fever syndromes carry mutations in *NLRP12*. More than 20 *NLRP12* mutations have been reported (*27–37*). Many of these mutations have been reported to enhance NF-κB signaling (*27, 30, 31, 33*), however, whether they cause caspase-1-depenent cleavage of IL-1β has not been investigated. These patients carry NLRP12 mutations that cause single amino acid substitutions or truncations due to either premature stop codons or reading frame shifts (Fig. S3B). To test their effect on IL-1β cleavage, we generated the corresponding mutations in mouse NLRP12 (Fig. 3B). All the mutants expressed well in HEK293T/17 cells, with similar levels of expression (Fig. S3C). Notably, though autoactivation was not observed in WT NLRP12-expressing cells, truncation mutants Q284X and W574X resulted in IL-1β cleavage (Fig. 3C). The Q284X mutant, which only contains the PYD and FISNA domains, appeared to have the strongest autoactivation, as demonstrated by the robust cleavage of IL-1β. In comparison to the truncation mutants, the reading frameshift mutants V628T-fs and R731fs, which both truncate the LRR, resulted in modest IL-1β cleavage (Fig. 3C). These results are consistent with the basic biochemistry common to the NLR family, which are often activated by truncation mutations. Most interestingly, the L551R mutant also resulted in strong IL-1β cleavage (Fig. 3C); this was the only full-length protein for which we observed autoactivation.

NLRP12 protein sequence alignment analysis showed that amino acid residues 284, 551, and 574 were highly conserved among mammals (Fig. 3D). The other patient-associated mutation sites were also conserved (Fig. S3D). Although the exact 3D structure of NLRP12 has not been reported, structural predictions by AlphaFold are available. Leu^551^ is buried within the protein (Fig. S3E), suggesting that an arginine substitution could be detrimental.

These results support the conclusion that NLRP12 forms an inflammasome whose activation is strictly regulated, but can be activated by certain mutations.

### NLRP3 and NLRP12 have distinct expression profiles in myeloid cell subtypes

It is well established that NLRP3 is highly expressed in monocytes and macrophages. However, the expression pattern of NLRP12 is less defined. ImmGen, BioGPS, and the Mouse Cell Atlas all indicate that *Nlrp12* is mainly expressed in granulocytes, especially neutrophils and eosinophils, but is poorly expressed in other immune cells, including monocytes, macrophages, and dendritic cells (Fig. S 4A, S4B, S4C). We performed qPCR from mouse tissues and found that *Nlrp12* was highly expressed in bone marrow and white blood cells (Fig. 4A). RNA-seq data demonstrated that the expression divergence of *Nlrp3* and *Nlrp12* occurs during hematopoietic stem cell (HSCs) differentiation through the myeloid lineage. Monocyte-dendritic cell progenitor (MDP)-derived cells (including monocyte-committed progenitors [cMoP], and the monocytes [M-mono] that they produce) express very low *Nlrp12*. In contrast, *Nlrp12* expression increases as granulocyte-monocyte progenitors (GMP) differentiate to their lineage-committed progeny, granulocyte progenitors (GP), which produce neutrophils. The branch towards GMP-derived monocyte progenitors (MP), also expressed *Nlrp12*, however this expression diminished upon differentiation into monocytes (G-mono) (Fig. 4B and Fig. S4D). In contrast, *Nlrp3* expression increases during monocyte differentiation from either GMPs or MDPs, but *Nlrp3* remains very low in GP (Fig. 4B and Fig. S4D). There were higher levels in the MDP pathway compared to the GMP pathway at each differentiation stage, however, both have high expression so slightly higher expression may not have a functional role. Note that terminally differentiated neutrophils and eosinophils were not evaluated in this experiment. Taken together, the summation of data indicates that *Nlrp12* expression is highly specific to neutrophils and eosinophils.

**Fig. 4.**
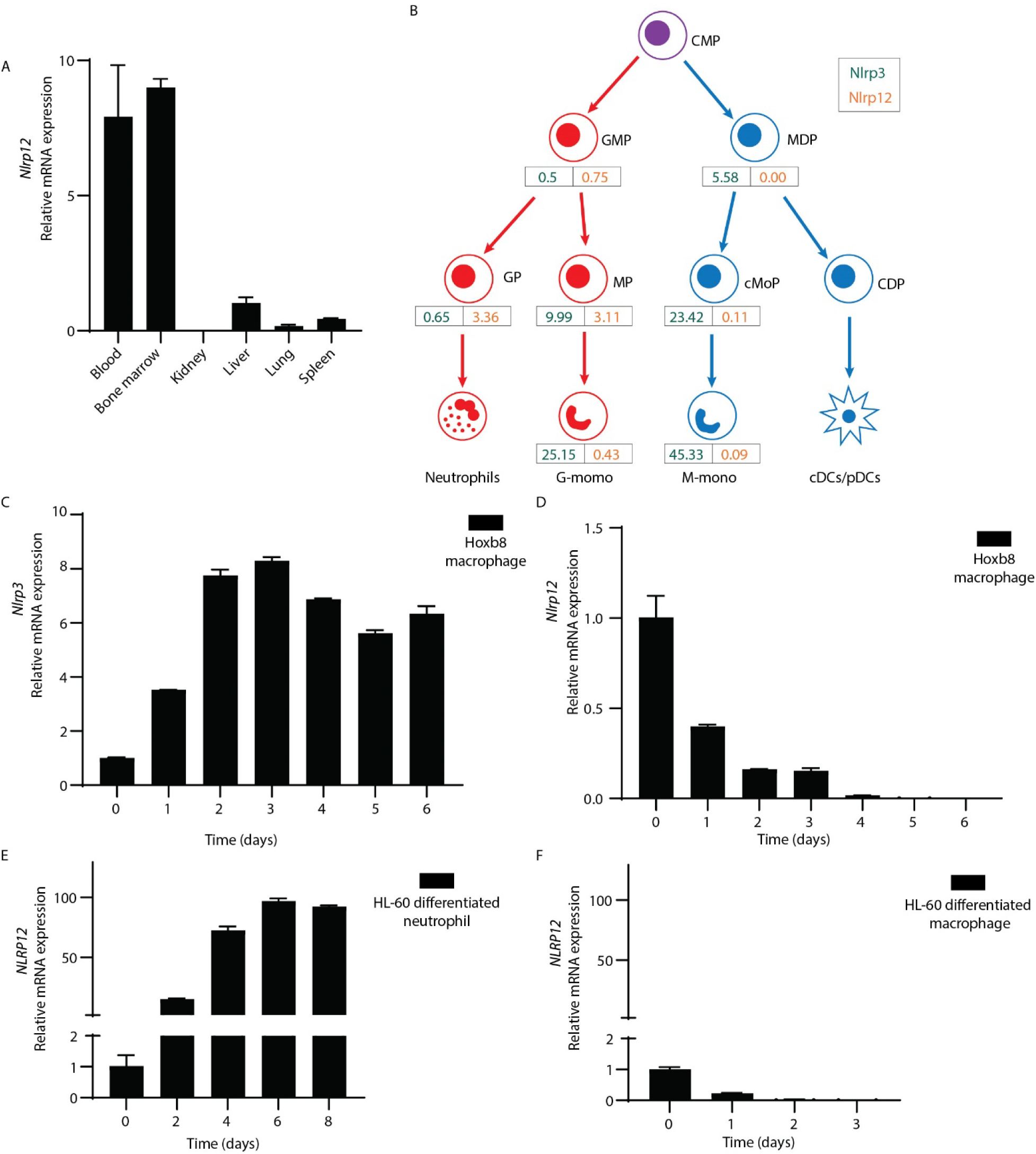
Specific expression profile of NLRP12 and NLRP3. (A) qRT-PCR analysis of *Nlrp12* expression in indicated mouse tissues. (B) *Nlrp3* (green) and *Nlrp12* (orange) expression by hematopoietic progenitors and monocytes was assessed in a previously published RNAseq dataset. RPKM values shown are means of duplicate samples of cells pooled from 20 mice each. CMP – common myeloid progenitors, GMP – granulocyte-monocyte progenitors, MDP – monocyte-dendritic cell progenitors, GP – granulocyte-committed progenitors, MP and cMoP – monocyte-committed progenitors, CDP – common dendritic cell progenitors, G-mono – GMP-derived monocytes, M-mono – MDP-derived monocytes, cDC and pDC – conventional and plasmacytoid dendritic cells. (C and D) qRT-PCR analysis of mouse (C) *Nlrp3* or (D) *Nlrp12* expression in Hoxb8-transduced progenitors that were induced to macrophage differentiation by L929 medium. (E and F) qRT-PCR analysis of *NLRP12* expression during HL-60 cell was induced to neutrophil-like cell differentiation by DMSO (1.25%) plus ATRA (1 μM) treatment or (F) macrophage differentiation by PMA (20 nM) treatment. All the results are representative of at least 3 independent experiments.

To study NLRP3 and NLRP12 during differentiation of macrophages and neutrophils in vitro, we transduced ER-Hoxb8 into common myeloid progenitor cells to create immortalized progenitors (*38*). We subjected these progenitor cells to macrophage differentiation with L929-conditioned media that contains M-CSF, and analyzed *Nlrp3* and *Nlrp12* expression by qPCR. Consistent with the above results, *Nlrp3* expression increased markedly during macrophage differentiation (Fig. 4C). On the contrary, the expression of *Nlrp12* dramatically decreased (Fig. 4D).

We also investigated expression of human NLRP12 using the HL-60 cell line derived from human acute promyelocytic leukemia (*39*). These cells can differentiate into either neutrophil-like or macrophage-like cells after different stimulation. Upon treatment with dimethyl sulfoxide (DMSO) and all-trans retinoic acid (ATRA) for 6 days, HL-60 cells differentiate into neutrophil-like cells, whereas upon PMA treatment for 3 days, HL-60 cells differentiate into macrophages (*40*). qPCR results showed that *NLRP12* expression was significantly increased during neutrophil differentiation (Fig. 4E), but was dramatically decreased during macrophage differentiation (Fig. 4F). In contrast, *NLRP3* expression did not change a lot during differentiation (Fig. S4E and S4F). This agrees with the data from ImmGen, where *Nlrp3* is expressed in both macrophages and neutrophils (Fig. S4G)

Therefore, NLRP12 is specifically expressed in neutrophils and eosinophils, but is not detectable in macrophages. In contrast, NLRP3 is highly expressed in monocyte/macrophages, and has low but appreciable expression in neutrophils.

### Inverse toxicity of NLRP3 and NLRP12 to macrophages and neutrophils

Molecular phylogeny analysis of all the human and mouse NLRP proteins showed that among NLRs, NLRP3 and NLRP12 are most closely related to each other (Fig. 5A). Notably, NLRP12, NLRP3, NLRP6 and NLRP10, all of which were confirmed to result in IL-1β cleavage by the PYD domain scanning, clustered together. The AlphaFold prediction of NLRP12 was similar to NLRP3, except the PYD domain (Fig. S5A). In the NLRP12 prediction, the PYD blocked the LRR domain, which may explain why NLRP12 shows lower propensity to auto-activate upon overexpression.

**Fig. 5.**
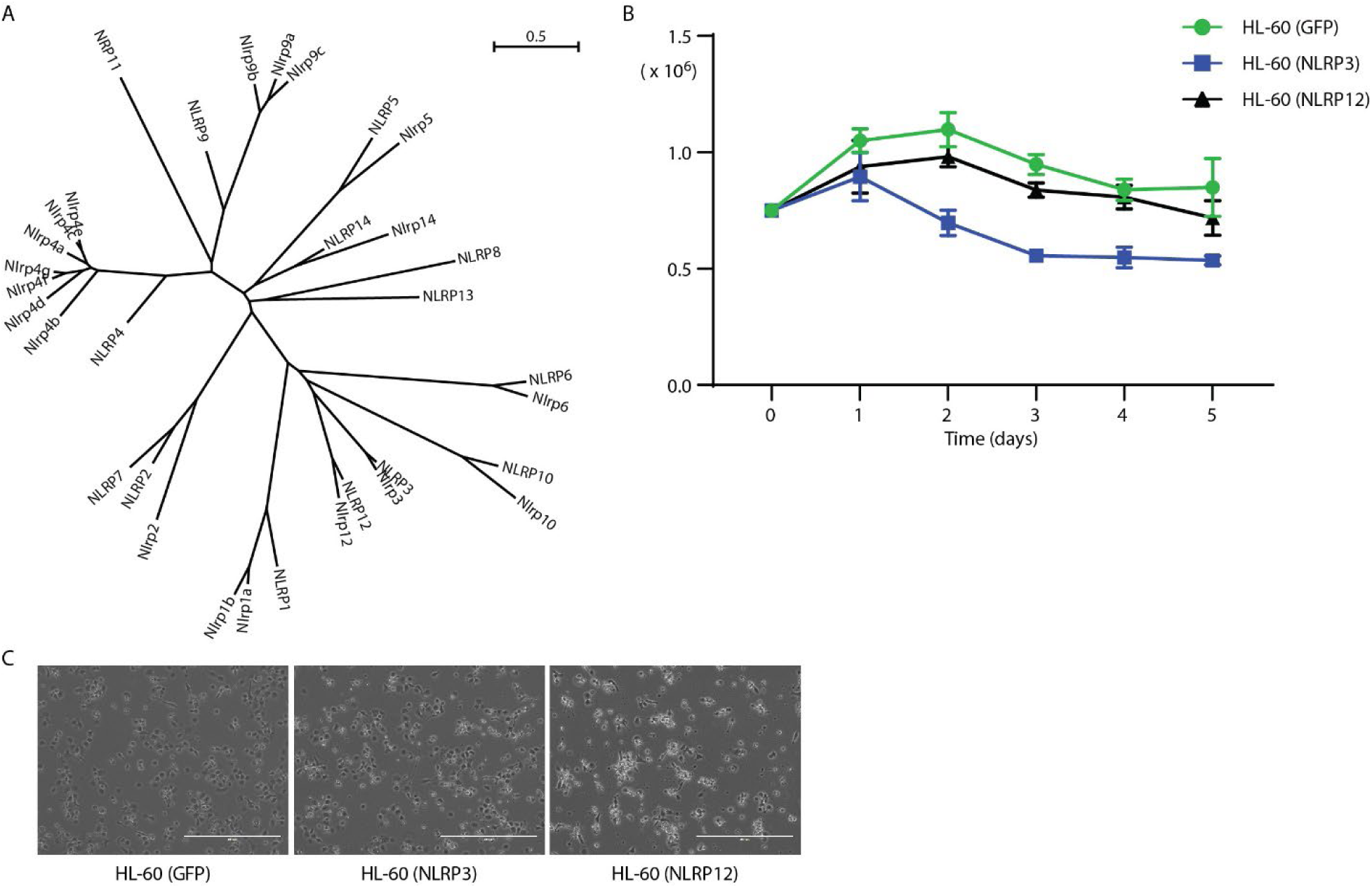
NLRP12 is toxic to macrophages. (A) Phylogenetic analysis of all the human and mouse NLRPs proteins. NLRP protein sequences were downloaded from NCBI. The sequences were analyzed by Seaview, which built the tree. (B and C) HL-60 cells were stably transduced with GFP, *NLRP3*, or *NLRP12*. (B) Stable cells were stimulated with DMSO (1.25%) plus ATRA (1 μM). The live cell numbers were counted at the indicated time point. (C) Stable cells were stimulated with PMA (20 nM) for 3 days and imaged by microcopy. Scale bars, 400μm. Results in (B) and (C) are representative of at least 3 independent experiments.

To study the role of the expression patterns during the differentiation of macrophages and neutrophils, we constructed HL-60 stable cell lines that ectopically-expressed mouse *Nlrp3* or *Nlrp12*. We subjected these cells to neutrophil differentiation using DMSO and ATRA as before and assessed viability by counting the total number of non-adherent, neutrophil-like cells. NLRP12-overexpressing cells showed similar surviving cell numbers compared to the GFP-transfected control (Fig. 5B). In contrast, over-expression of NLRP3 yielded fewer cells during the differentiation process beginning at day 2 (Fig. 5B). On the other hand, during PMA-induced macrophage differentiation, we imaged each well to estimate the number of successfully differentiated macrophage-like cells that were adherent in the well. There were similar numbers of NLRP3-expressing cells compared to the GFP-expressing control cells (Fig. 5C). In contrast, the number of NLRP12-expressing cells was markedly lower, and these cells had abnormal morphology (Fig. 5C). Together, these results suggest that high levels of NLRP3 expression may be incompatible with neutrophil differentiation and NLRP12 expression may be incompatible with monocyte/macrophage differentiation.

### Cytosolic Location of NLRP3 and NLRP12

To explore the subcellular location of NLRP3 and NLRP12, we constructed Hela stable cell lines which ectopically-expressed the wild type and mutant of *Nlrp3* and *Nlrp12*. In the resting state, both NLRP3(WT) and NLRP3(R258W) were located in the cytosol. But after nigericin stimulation, NLRP3(WT) formed multiple foci in a peri-nuclear region, which matched the previous report that NLRP3 can be recruited to dispersed trans-Golgi network (dTGN) upon stimulation (*41, 42*). The expression of NLRP3(R258W) was weaker than wild type, which was coincident with the previous immunoblotting result (Fig. 2A). Furthermore, the location was similar under resting state, which was also in the cytosol (Fig. 6). To our surprise, the mutant NLRP3(R258W) did not form puncta after stimulation like NLRP3(WT). In addition to HeLa cells, we also transduced COS-1 cells with *Nlrp3*. NLRP3(WT) was also located in the cytoplasm in rest state, and formed numerous foci after nigericin stimulation (Fig. S6B). Interestingly, there were already some foci before stimulation, which might be due to some specific co-factors that only existed or were rich in COS-1 cells.

**Fig. 6.**
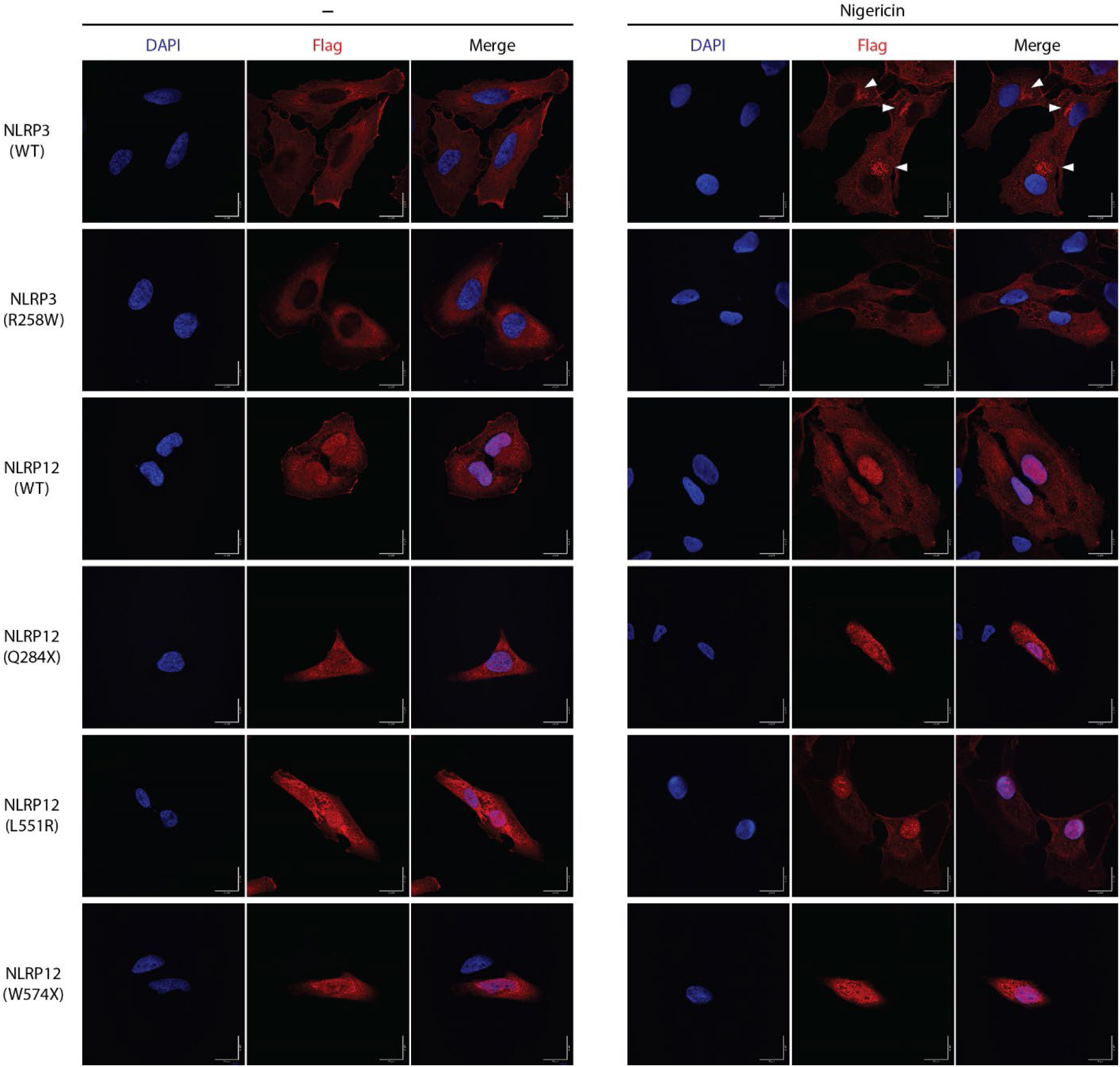
Subcellular location of NLRP3 and NLRP12. HeLa cells were stably transduced with mouse NLRP3 or NLRP12 and were stimulated with nigericin or not for 60 min. Immunostaining was performed for the Flag epitope tag and cells visualized by confocal microscopy. Triangles indicate NLRP3 foci. Scale bar, 20μm. All the images are representative of at least 3 independent experiments.

NLRP12(WT) was distributed not only in the cytosol but also was present in the nucleus (Fig. 6). NLRP12(WT) did not form foci after nigericin treatment, which is different from NLRP3(WT). The cellular location of activating point mutation NLRP12(L551R) and NLRP12(F402L) were similar to NLRP12(WT), regardless of nigericin treatment. The truncation (NLRP12(Q284X) and NLRP12(W574X)) and frame shift (NLRP12 (V628T-fs) and NLRP12(R731-fs)) mutations were distributed throughout the cell, and did not response to nigericin stimulation (Fig. 6 and Fig. S6A). In COS-1 cells, NLRP12(WT) showed less nuclear location regardless of nigericin treatment, and again did not form foci after nigericin stimulation. The slight differences in nuclear exclusion may be due to cell line specific properties and suggest that the nuclear localization may not reflect an important aspect of NLRP12 function, but rather may reflect the ability of each cell line to exclude various proteins from the nucleus. The different localization patterns of NLRP3 and NLRP12 suggest that the activation mechanism of NLRP12 could be distinct from that of NLRP3.

## Discussion

Both inflammation and pyroptosis are double-edged swords, which are necessary to battle against pathogens but can also cause damage to tissues. Therefore, inflammasome activation should be under tight control. Here, we demonstrate that the NLRP proteins NLRP6 and NLRP12 can also form inflammasomes, similar to NLRP3. However, these three inflammasomes are expressed in distinct cell types. The NLRP6 inflammasome is specifically expressed in IECs, while the NLRP12 inflammasome specifically exists in neutrophils and eosinophils. Meanwhile, NLRP3 is expressed in more cell types, including most myeloid cells such as macrophages, monocytes, DCs, mast cells, and neutrophils. Similarly, recent reports showed that NLRP10 also has restricted expression to either keratinocytes in the skin or intestinal epithelial cells of the distal colon (*18, 19*). All these cell types are exposed to infection by pathogens, but the different expression profiles of NLRP3, NLRP6, NLRP10, and NLRP12 probably reflects that each cell type has a distinct cell physiology that pathogens would evolve to target.

Each NLR could monitor a different aspect of cellular physiology or morphology that is unique to a specific cell type, and activate in response to a pathogen perturbing that aspect of cellular biology. To illustrate this concept, we will speculate on some possible examples of cell type-specific functions that could be monitored by these NLRs. NLRP6 is specifically expressed in IECs, which have vastly different morphology and immunologic capabilities compared to macrophages and neutrophils (*43*). IECs form a barrier and must absorb nutrients from the gut lumen (*43, 44*). To accomplish this, IECs have a large surface area created by microvilli, which are formed around densely polymerized actin structures (*45*). Microvilli are not present in immune cells. Speculatively, NLRP6 could function as a sensor (or a guard) (*46*) that monitors for the integrity of microvilli. Thus, if a pathogen disrupted the microvilli, this could cause NLRP6 to activate. Such a hypothetical case or another cell type specific property could explain why NLRP6 is highly autoactivate in HEK293T/17 cells – perhaps it is activating in response to the absence of microvilli. One would then expect NLRP6 to detect enteropathogenic *Escherichia coli* or *Citrobacter rodentium*, which remodel the microvilli; however, as these are successful in their native hosts, the pathogens would be expected to use virulence factors to evade NLRP6 detection. Interestingly the two other organs that express NLRP6 are the liver and kidney, both of which contain epithelial cells with microvilli (*45*). Hepatocytes in the liver use microvilli to filter the blood, whereas brush border cells in the proximal tubule of the kidney use microvilli in ion exchange in the formation of urine.

NLRP12 expression is highly restricted to neutrophils and eosinophils. Neutrophils share many properties with macrophages; however, our data suggest that NLRP12 might autoactivate in macrophages. There are many differences between macrophages and neutrophils. Neutrophils contain pre-formed granules that are essential for their antimicrobial function, which is also a characteristic of eosinophils (*47*). These granules must fuse with the phagosome in neutrophils, and also can be degranulated in neutrophils and eosinophils. Neutrophils and eosinophils also make considerably more reactive oxygen species using the NADPH oxidase than macrophages are capable of producing (*47*). We speculate that one of these properties could be a function that NLRP12 monitors, and activates only when the function is impeded by pathogen effectors.

Previous research reported that NLRP12 negatively regulates the NF-κB pathway via affecting the stability of NF-κB inducing kinase (NIK) (*48*). There were also reports showing that NLRP12 was a tumor suppressor gene. Allen. *et al* found *Nlrp12^−/−^*mice were highly susceptible to colitis and colitis-associated colon cancer (*49, 50*). The authors attributed this to multiple signaling pathways, especially non-canonical NF-κB, all of which were negatively regulated by NLRP12. Udden *et al.* reported that *Nlrp12^−/−^*mice were highly susceptible to diethylnitrosamine-induced hepatocellular carcinoma (HCC) (*51*). The authors implicated NLRP12 as a negative regulator downregulation of JNK-dependent inflammation and proliferation of hepatocytes (*51*). Ulland *et al* reported that in C57/B6J mouse, a missense mutation of *Nlrp12* (R1034K) caused neutrophil recruitment defect (*52*), which has been reported to affect tumorigenesis. Further investigation of these phenotypes could benefit from the knowledge that NLRP12 is an inflammasome, and that it is specifically expressed in neutrophils and eosinophils.

Our data support the conclusion of other prior studies that NLRP12 can form inflammasomes. Vladimer *et al.* found that NLRP12 recognized infection by a modified strain of *Yersinia pestis*, and mediated activation of caspase-1 and release of IL-1β and IL-18 (*53*). However, although the researchers had mentioned that *Nlrp12* was expressed more in neutrophils than macrophages (*53*), this conclusion has not been widely appreciated in subsequent papers. Here, we reinforce the conclusions of Vladimer *et al.* by showing that NLRP12 is primarily expressed in neutrophils, and also in eosinophils. Recent work from Coombs et al. used PBMCs stimulated with LPS for 24 hours, which should not contain neutrophils or eosinophils (*54*). Other recent work from Sundaram *et al.* studied NLRP12 primarily in murine bone marrow-derived macrophages, a cell type that we found to express undetectable levels of *Nlrp12* message (*55*). The researchers reported NLRP12 activated in response to heme and mediated PANoptosis with ASC, caspase-8, and RIPK3 (*55*). In that paper, the researchers found the expression of *Nlrp12* increased more than 10 times after 36 hours of LPS or Pam_3_CSK_4_ stimulation (*55*), which is longer than typical priming treatments used during in vitro studies. Similarly, the pyrin inflammasome is normally expressed by neutrophils, and to a lesser extent monocytes, but not by macrophages. However, 24 hours of TLR or cytokine stimulation will modestly induce pyrin expression in monocytes or macrophages (*56, 57*). Thus, prolonged stimulation in macrophages can induce expression of the NLRP12 or pyrin inflammasomes, which could be relevant to in vivo infections where pathogens typically linger for many days.

Clinical researchers have reported an autoinflammatory disease caused by NLRP12 mutation (called NLRP12-AID). The patients typically presented with periodic fever, urticaria-like rash, arthralgia/arthritis, myalgia, and lymphadenopathy (*31*). Among the patients, most developed the disease in childhood. Through exon sequence or other methods, the researchers have reported more than 20 NLRP12 variants, including point mutation, truncation, and reading frame shifts. Some researchers have found increased inflammatory cytokine secretion, including IL-1β, TNFα, and IFN-γ in some patients’ serum (*27, 29, 33*). The patients’ PBMC cells were more sensitive to LPS stimulation than control cells from health people. The authors attributed these to activation of the NF-κB pathway; a few mutants tested with NF-κB responsive luciferase showed a loss of the inhibitory efforts of NLRP12 (*30, 31*), but many mutants did not show this activity. A recent report suggested that NLRP12 is not an inflammasome, but instead is a tonic inhibitor of NLRP3, and that the patient-associated mutations did not activate ASC signaling, including the human V635T frame shift that deletes the LRR domain (*54*). In direct contrast to this, we now show that several mutations in NLRP12-AID patients (including the mouse equivalent of V635T frame shift) cause spontaneous activation of caspase-1. However, other NLRP12 mutations did not cause caspase-1 activation. We speculate that some of these patient mutations could cause spontaneous activity in neutrophils, and that there may be neutrophil specific factors that are missing from HEK293T/17 cells that cause some of these mutations to fail to autoactivate in our studies. This could be analogous to how NLRP3 needs NEK7; NLRP12 may require a co-factor that is only expressed in neutrophils. Alternately, it may be that some NLRP12 amino acid substitutions are not actually causative of the patient’s disease. The F402L mutation has been reported by several clinical papers to be present in patients with autoinflammatory syndromes, and F402 is highly conserved among species (Fig. S 3D). However, F402L is a common polymorphism found in humans who do not have autoinflammatory disease, and thus this mutation has been proposed to not be causative of autoinflammatory disease (*28*), a conclusion that is supported by our data where F402L is not an activating mutation.

IL-1β blockade has been used in two NLRP12-AID patients – Anakinra therapy achieved a marked clinical improvement at the first two weeks, secretion of IL-1β by peripheral blood mononuclear cells (PBMCs) decreased significantly within two months of treatment (*58*). However, over time the efficacy was lost and treatment was discontinued after 14 months(*58*). It may be that these patients have symptoms driven by the combined effects of IL-1β and IL-18. In this regard, NLRC4 auto-activating mutations received therapeutic benefit from IL-18 blockade (*59*). However, pyroptosis also releases multiple cytosolic molecules that are inflammatory, and patients might additionally need to be treated with drugs that inhibit GSDMD (*60*).

## Material and Methods

### Antibodies and Reagents

The following antibodies were used: mouse monoclonal anti-FLAG M2 antibody (F1804, SIGMA); mouse monoclonal anti-HA (16B12) (MMS-101P, Covance); myc antibody(9E10) (sc-40, Santa Cruz); Peroxidase-conjugated AffiniPure Goat Anti-Mouse IgG (H+L) (115-035-062, Jackson ImmunoResearch); Goat anti-Mouse IgG (H+L) Highly Cross-Adsorbed Secondary Antibody, Alexa Fluor™ Plus 555 (A32727, Invitrogen).

The following reagents were used: AP20187 (SML2838, SIGMA); Nigericin (tlrl-nig-5, InvivoGen); TRIzol reagent (15596026, Invitrogen); SuperScript™ II Reverse Transcriptase (18064014, Invitrogen); RNaseOUT™ Recombinant Ribonuclease Inhibitor (10777019, Invitrogen); Deoxynucleotide (dNTP) Solution Mix (N0447L, NEB); Dimethyl sulfoxide (DMSO) (D4540, SIGMA); Retinoic acid (ATRA) (R2625, SIGMA); Phorbol 12-myristate 13-acetate (PMA) (tlrl-pma, InvivoGen); (Z)-4-Hydroxytamoxifen (4-OHT) (74052, STEMCELL); Recombinant Murine GM-CSF (315-03, Peprotech); Phenylmethanesulfonyl fluoride solution (PMSF) (93482, SIGMA); Protease Inhibitor Cocktail (HY-K0010, MCE); Puromycin (P8833, SIGMA); RNAlater Stabilization Solution(AM7020, Invitrogen); PowerTrack SYBR Green Master Mix (A46110, Invitrogen); Polybrene (H9268, SIGMA); Paraformaldehyde (PFA) (P6148, SIGMA); DAPI (D8417, SIGMA).

### Cell culture

HEK293T/17 (CRL-11268, ATCC), HeLa (CCL-2, ATCC), COS-1 (CRL-1650, ATCC) and L-929 (CCL-1, ATCC) cells were maintained in DMEM (11995073, Gibco) supplemented with 1% Penicillin-Streptomycin (15140122, Gibco) and 10% Fetal Bovine Serum (SH30396.03, Cytiva). HL-60 (CCL-240, ATCC) cell was maintained in RPMI1640 (11875093, Gibco) supplemented with 1% Penicillin-Streptomycin (15140122, Gibco) and 10% Fetal Bovine Serum (10082147, GIBCO). For HL-60 cell neutrophil differentiation, cells were induced with a final concentration of 1.25% DMSO plus 1 μM ATRA for 6 days. For HL-60 cell macrophage differentiation, cells were induced with 20 nM PMA for at least 3 days. Hoxb8 progenitor cells were maintained in RPMI1640 (11875093, Gibco) supplemented with 1% Penicillin-Streptomycin (15140122, Gibco), 20 ng/mL, GM-CSF, 100 nM 4-OHT and 10% Fetal Bovine Serum (10082147, GIBCO). For Hoxb8 progenitor cell differentiation, cells were induced by withdrawing 4-OHT or L-cell medium, which contained DMEM, 10% L-929 cell culture medium and 10% Fetal Bovine Serum (SH30396.03, Cytiva). All the cell lines were tested regularly for mycoplasma infection by PCR.

### qRT-PCR

About 20ug tissue samples or more than 4 x 10^6^ cells were harvested from C57/B6J mice for RNA extraction with TRIzol reagent. For tissue samples, homogenization is essential. 1ug total RNA was subjected to reverse transcription through SuperScript II Reverse Transcriptase. The gene expression was assayed by normal SYBR green method with PowerTrack SYBR Green Master Mix. The results were analyzed by ΔΔCT method. The mouse tissue samples can be stored in RNAlater Stabilization Solution at -80°C if the experiments were not processed immediately. The cDNA samples can be stored at -80°C, too.

The PCR primers were designed by Primer-Blast (https://www.ncbi.nlm.nih.gov/tools/primer-blast/), sequence as follow: mouse 18S rRNA (forward: GGCCGTTCTTAGTTGGTGGA, reverse: TCAATCTCGGGTGGCTGAAC); mouse Gapdh (forward: GAAGGTCGGTGTGAACGGAT, reverse: TTCCCATTCTCGGCCTTGAC); *mouse Nlrp6* (forward: AGCTGTAGAAATGACCCGGC, reverse: GAACGCTGACACGGAGAGAA); mouse Nlrp3 (forward: AGAGTGGATGGGTTTGCTGG, reverse: CGTGTAGCGACTGTTGAGGT); mouse Nlrp12(forward: TGGCTCTCAGCACCTTTCAG, reverse: AGAGACATCCAAAGGGCACG); human GAPDH (forward: GGAAGGTGAAGGTCGGAGTC, reverse: TGGAATTTGCCATGGGTGGA): human NLRP6 (forward: ACCACAAAACAACTGCCAGC, reverse: CCTCAGGGCCTCAGAAAGGT); human NLRP3 (forward: CACTGTCCCTGGGGTTTCTC, reverse: CCCGGCAAAAACTGGAAGTG); human NLRP12 (forward: TGTGGGAGAGAGGACAGAGAG, reverse: AGGTTTCCTGGGGATCTTTTCT).

### Transfection and Immunoblotting analysis

The HEK293T/17 cells were transiently transfected with Calcium Phosphate Transfection method. The reagents are 2.5 M CaCl_2_ and 2x HEPES buffer (50 mM HEPES, pH 7.05, 280 mM NaCl, 1.5 mM Na_2_PO_4_). Medium may be changed with fresh medium 8 hours post transfection. The cells were harvested with DPBS (14190144, Gibco), and lysed with RIPA buffer (50 mM Tris, pH 7.4, 150 mM NaCl, 1% Triton, 0.1% sodium deoxycholate, 0.01% SDS, 1 mM EDTA, 1 mM EGTA, 2 mM NaF, 1 mM Na3VO4, 1 mM β-glycerophosphate, 1 mM PMSF, protease inhibitor cocktail). The Cell lysates were subjected to western blotting with indicated antibody.

### Stable cell line construction

The cDNA of *Nlrp3* and *Nlrp12* were subcloned into lentiviral expression vector pLenti-EF1a-C-Myc-DDK-IRES-Puro (a gift from Dr. Youssef Aachoui). Lentiviral package process was followed the addgene protocol (https://www.addgene.org/protocols/lentivirus-production/). Briefly, the expression plasmid, together with viral package plasmid psPAX2 and pMD2.G, were transfected into HEK293T/17 cells at the ration of 4:3:1. Medium that contains lentiviral particles, was harvested 2 days later, and filtered by 0.45μM low protein binding filter. Hela, COS-1 and HL-60 cells were transduced with lentiviral particles in the presence of polybrene (5μg/mL). The transduced cells were selected by puromycin(2μg/mL) 2 days later for 1 week. The protein expression was detected by immunoblotting.

### AlphaFold prediction

The structure of mouse NLRP3 and NLRP12 were from AlphaFold database; the peptide structures were predicted via colabFolad notebook (*61, 62*).

### RNAseq analysis

Analysis of Nlrp3 and Nlrp12 expression by hematopoietic progenitors and monocytes was performed using a published RNAseq dataset (GEO: GSE88982) of cells isolated from mouse bone marrow (*63*)(https://pubmed.ncbi.nlm.nih.gov/29166589/). Ex vivo GMP, GP and MDP, as well as monocyte-committed progenitors and classical monocytes derived from GMP and MDP in vitro, were analyzed by RNAseq.

### Immunofluorescence

Cells were plated on circular cover glasses (12-541-001, Fisherbrand) in 24 well plates. After 24 hours, the cells were treated with Nigericin (20 μM) for 60 min or not. Then, cells were processed as follow: fixed by 4% PFA (pH 7.4 in DPBS) for 15 min; permeabilized by 0.25% Triton X-100 in PBS for 15 min; blocked by blocking buffer (5% normal goat serum in PBS-0.05% Tween 20(PBS-T)) for 1hour; immunostained overnight with indicated primary antibodies in a humidified chamber at 4 °C; washed 3 times with PBS-T; subsequently incubated with Alexa Fluor Plus 555-conjugated secondary antibodies for 1 h; washed 3 times again and stained with DAPI (10 μg/mL) for 10 min; Mounted on the slide with anti-fade mounting medium. Images were captured on Zeiss LSM 780, and analysis by ImageJ (Fiji).

## Acknowledgments

The authors thank Dr. Youssef Aachoui from UAMS for pLenti-EF1a-C-Myc-DDK-IRES-Puro, Dr. Hasan Zaki from UTSW for Nlrp12 expression plasmid. HSG is supported by NIH grant R01 AI134987.

## Author contributions

Conceptualization: B.W., E.A.M.

Methodology: B.W., E.A.M., H.S.G., K.N., Z.R.B.

Investigation: B.W.

Visualization: B.W.,

Funding acquisition: E.A.M., H.S.G.

Project administration: E.A.M., B.W.

Supervision: E.A.M.

Writing – original draft: B.W.

Writing – review & editing: E.A.M with all other authors

## Data and materials availability

All reagents and plasmids used in this study are available upon request. All data needed to evaluate the conclusions in the paper are present in the paper or the Supplementary Materials.

---

**Fig. S1.**
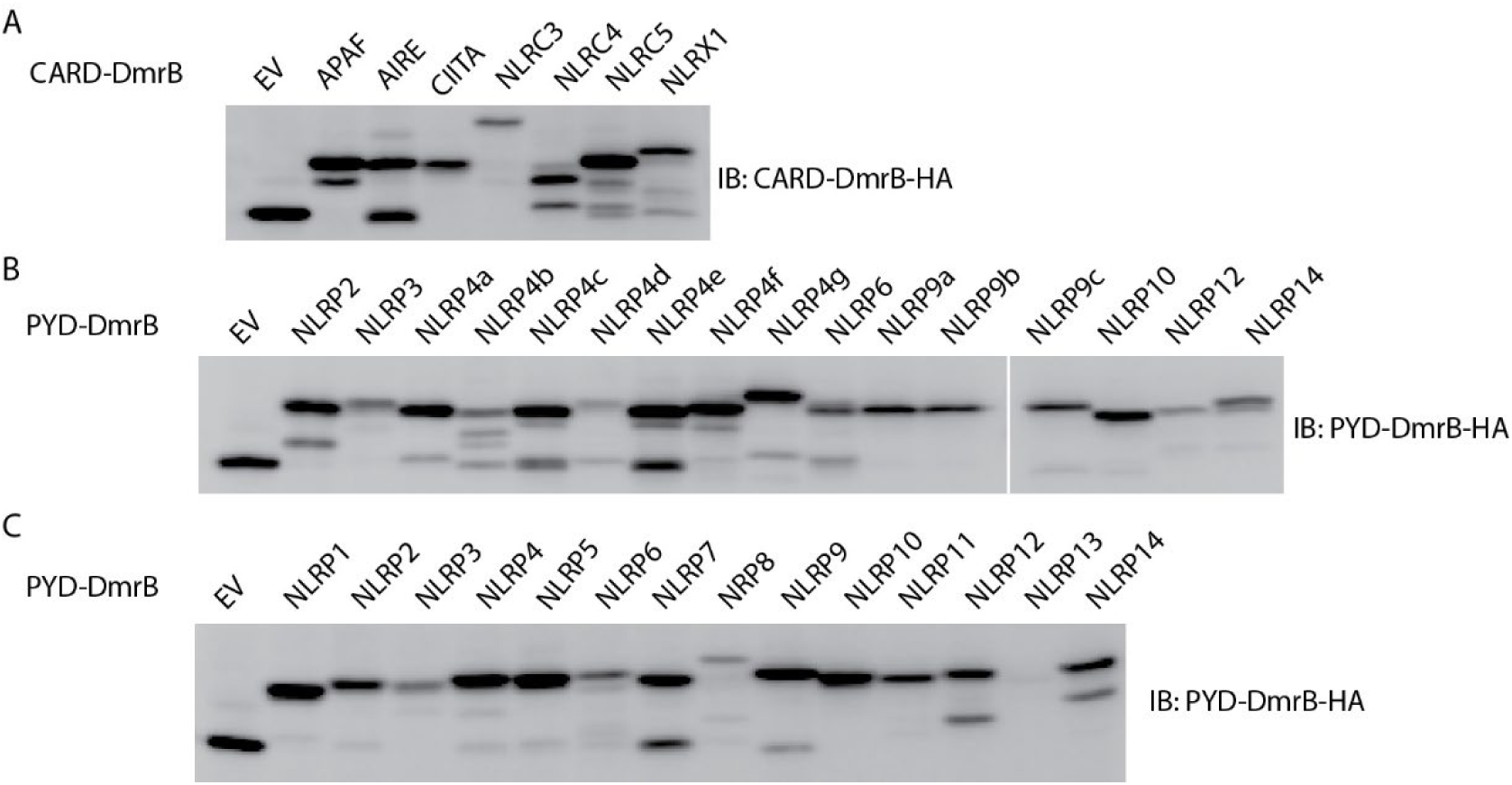
The expression of CARD and PYD domain fusion proteins. (A) The expression of mouse CARD domain fusion proteins. The CARD domain from *Apaf*, *Aire*, *Ciita*, *Nlrc3*, *Nlrc4*, and N-terminal domain of *Nlrx1* were subcloned and expressed together with tandem DmrB domain as fusion proteins. (B) The expression of mouse PYD domain fusion proteins. The PYD domain from *Nlrp2, Nlrp3, Nlrp4a*, *Nlrp4b*, *Nlrp4c*, *Nlrp4d*, *Nlrp4e*, *Nlrp4f*, *Nlrp4g, Nlrp6*, *Nlrp9a*, *Nlrp9b*, *Nlrp9c, Nlrp10*, *Nlrp12*, and *Nlrp14* were subcloned and expressed together with tandem DmrB domain as fusion proteins. (C) The expression of human PYD domain fusion proteins. The PYD domain from *NLRP1*, *NLRP2*, *NLRP3, NLRP4*, *NLRP5*, *NLRP6*, *NLRP7*, *NLRP8*, *NLRP9*, *NLRP10*, *NLRP11*, *NLRP12*, *NLRP13*, and *NLRP14* were subcloned and expressed together with tandem DmrB domain as fusion proteins. The HEK293T/17 cells were transiently transfected with indicated plasmid for 24h, and lysates were subjected to immunoblotting with indicated antibody. All the blotting results are representative of at least 3 independent experiments.

**Fig. S2.**
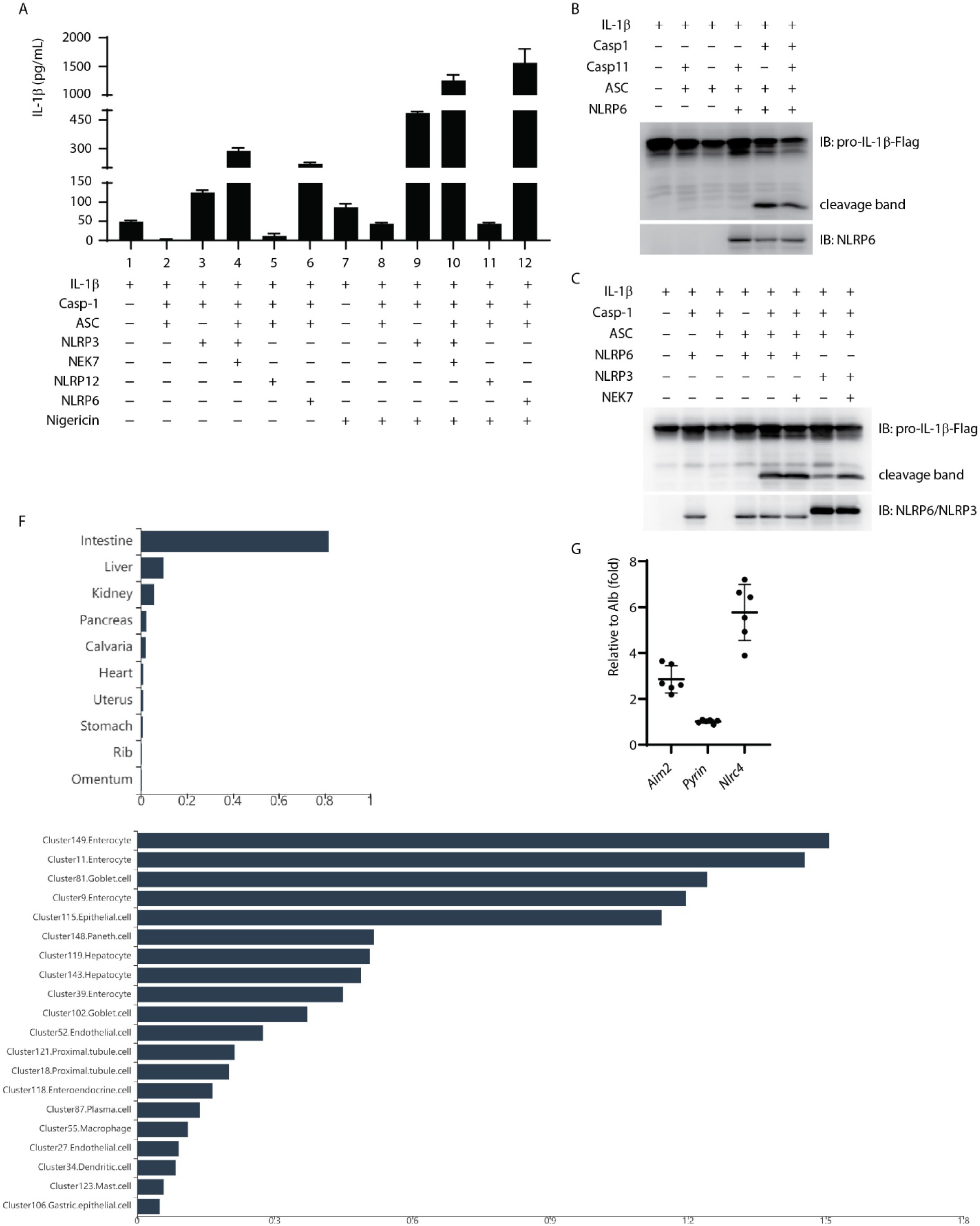
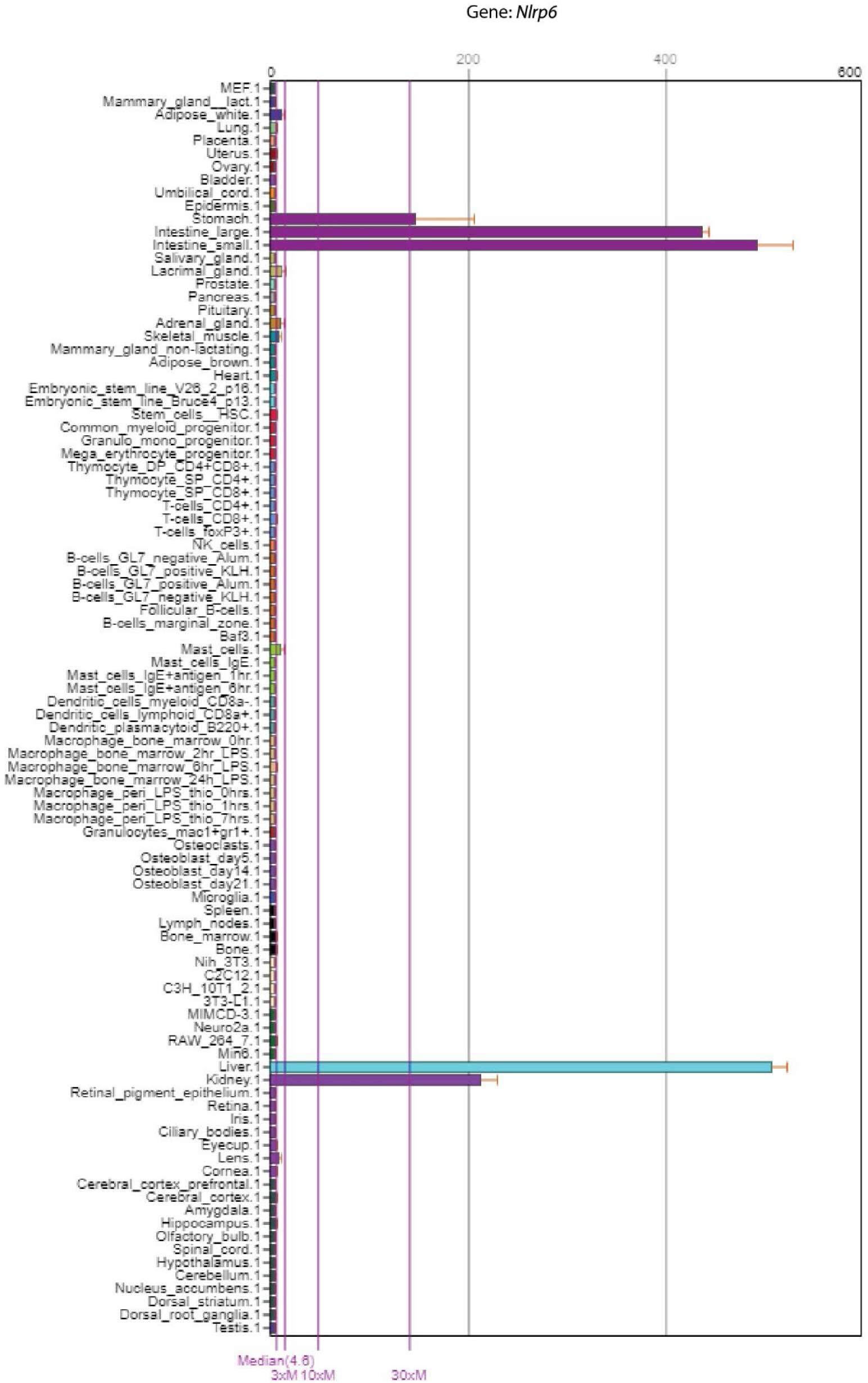
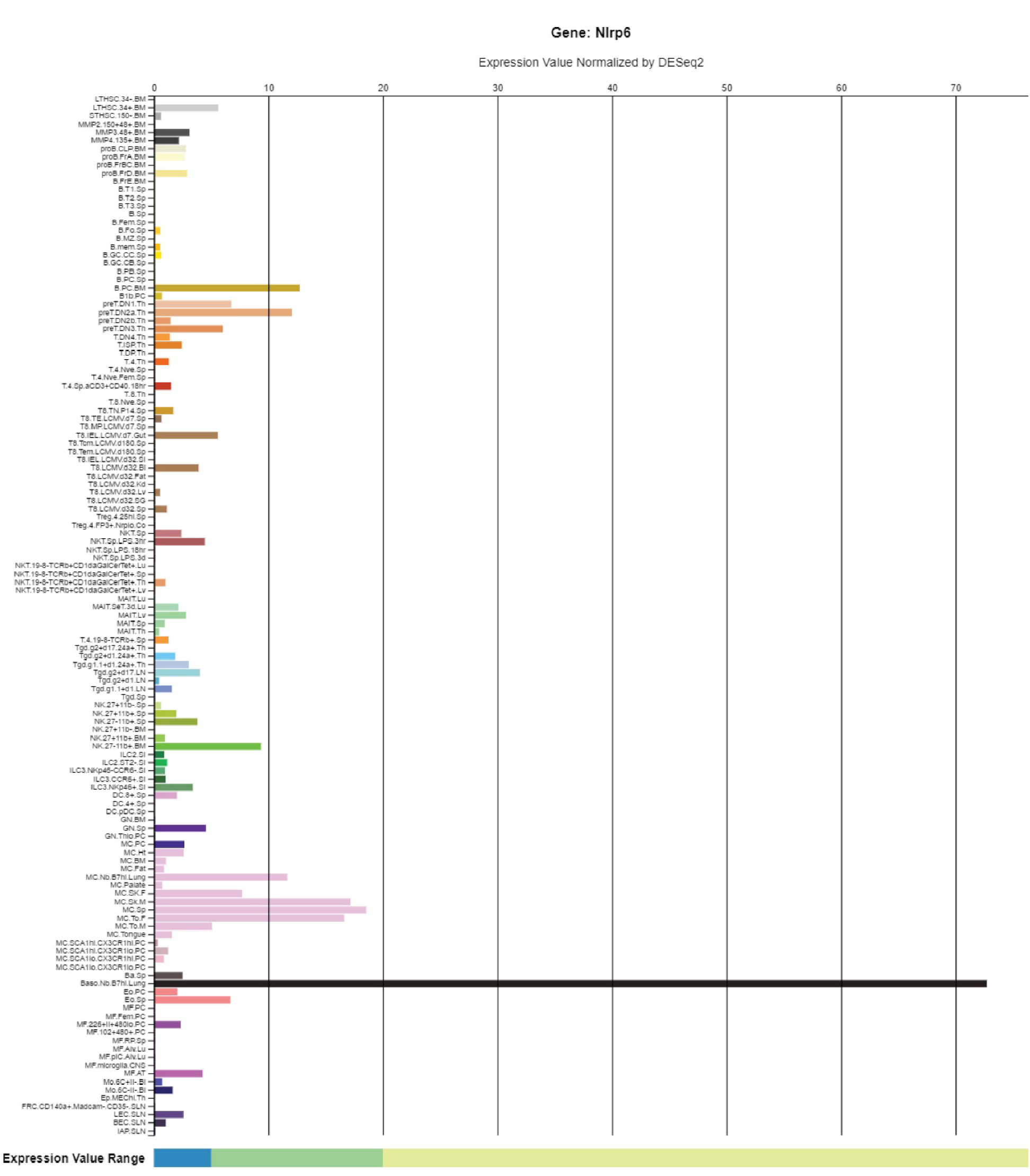
Reconstitution of inflammasome in vitro. (A) NLRP3, NLRP6 and NLRP12 inflammasome were reconstituted in HEK293T/17 cells using the indicated plasmids with or without nigericin treatment. IL-1β concentration in cell culture medium was assayed by ELISA. (B and C) Reconstitution of NLRP6 inflammasome with *Casp11* (B) and NEK7 (C) with indicated immunoblots. (D-F) The expression of mouse *Nlrp6* from BioGPS (D), ImmGen (E) and the Mouse Cell Altas (F). The results were downloaded from BioGPS, ImmGen (RNA-seq), and the Mouse Cell Altas websites. (G) The expression of *Aim2*, *Pyrin*, and *Nlrc4* in IECs. The original data is downloaded from NCBI (GDS3921). The results were normalized by albumin (*Alb*). The results in (A), (B) and (C) are representative of at least 3 independent experiments.

**Fig. S3.**
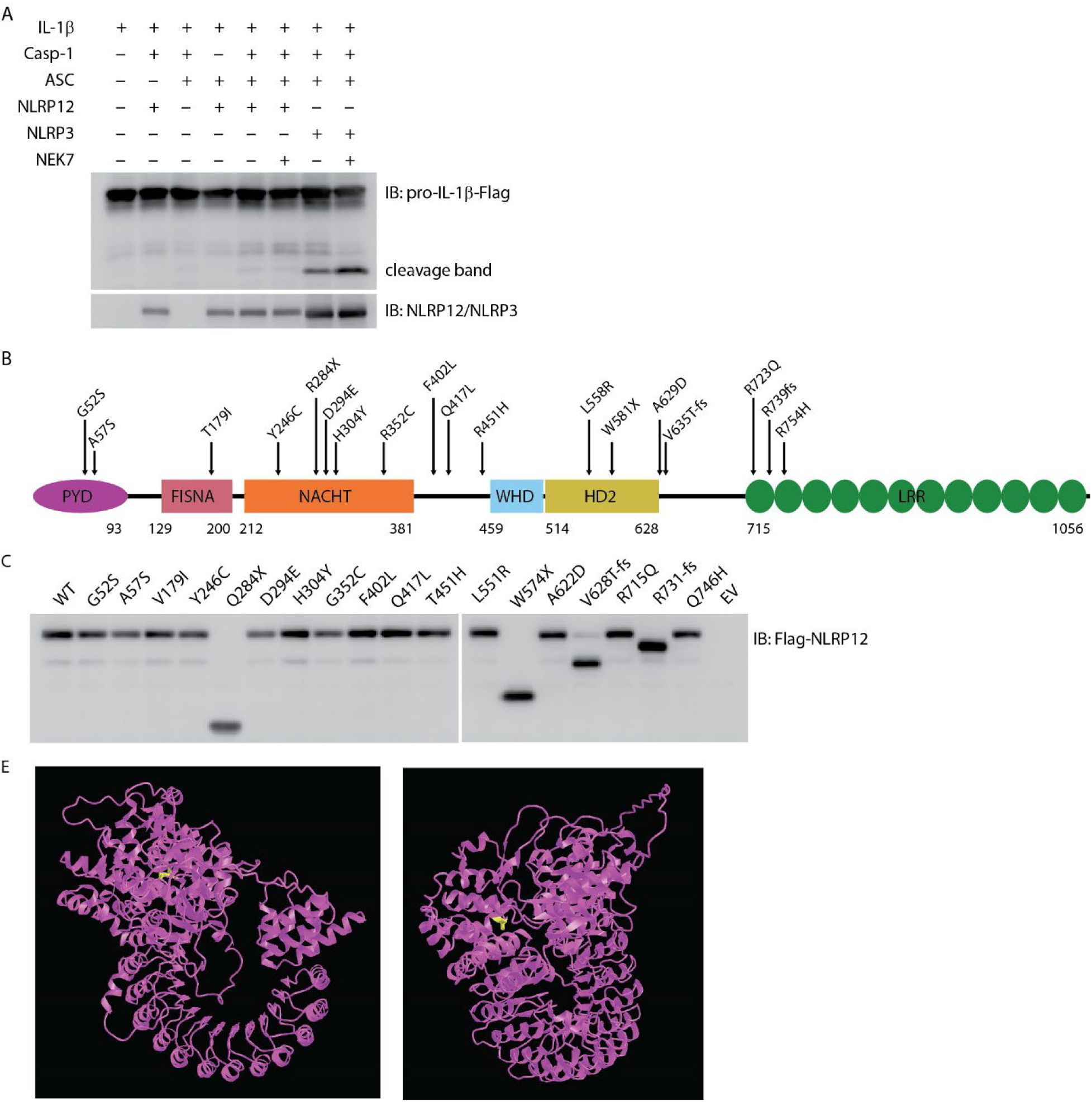
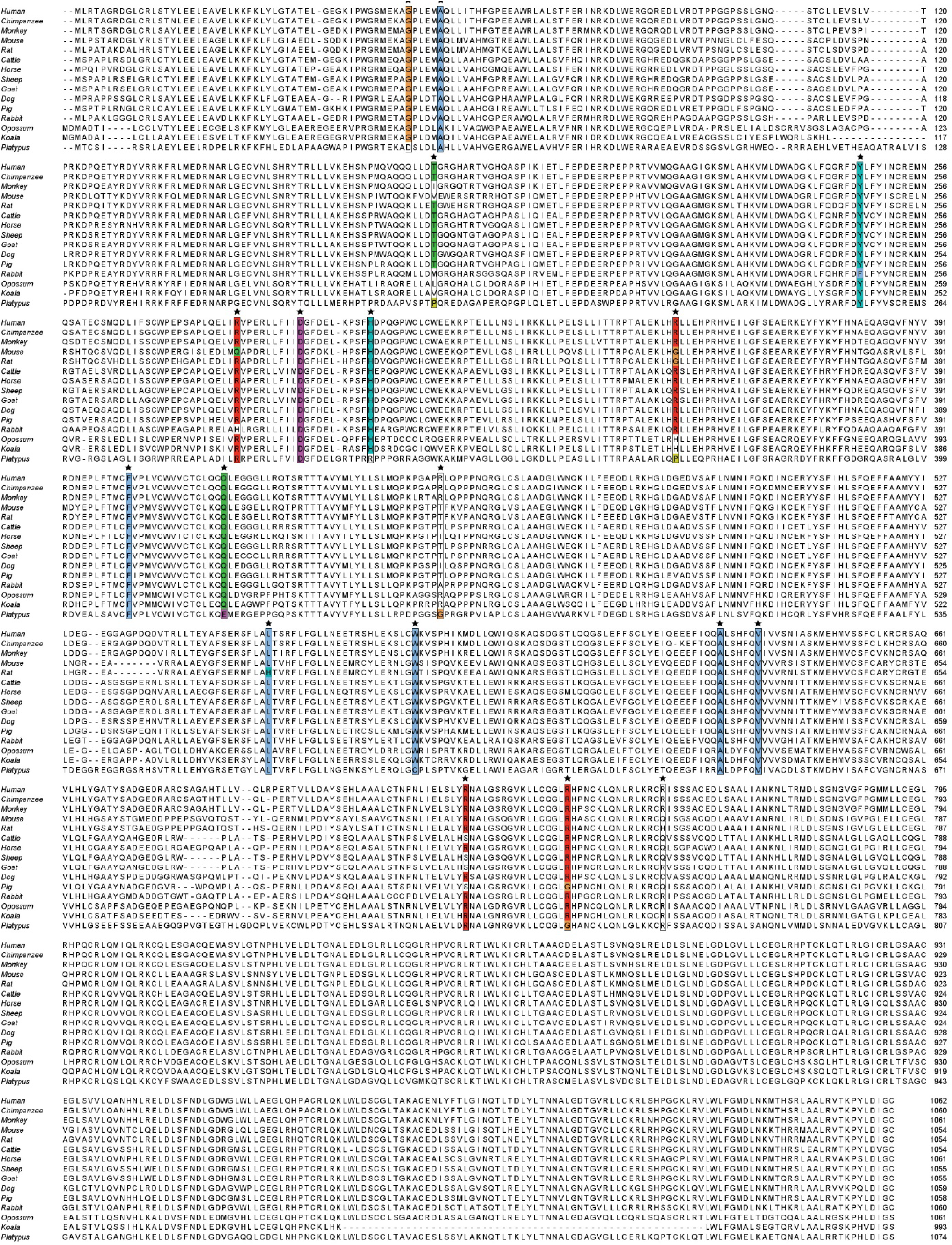
The character of NLRP12. (A) Reconstitution of NLRP12 inflammasome with IL-1β, Caspase-1, ASC, NEK7, NLRP12 (WT) and NLRP3 (as a positive control) with indicated immunoblots. (B) Schematic diagram of human NLRP12 mutants, which have been reported in human NLRP12-AID patients by clinical researchers before. (C) The expression of various NLRP12 mutants and wild type in HEK293T/17 cells. (D) Alignment of NLRP12 from indicated species. The asterisk indicated the mutation sites in Fig. S3B and Fig. 3B.(E) The predicted structure of mouse NLRP12 by AlphaFold. The date file was downloaded from AlphaFold protein structure database, and analyzed by iCn3D. (https://www.ncbi.nlm.nih.gov/Structure/icn3d/full.html). Leu^551^ was labeled yellow. The blotting results in (A) and (C) are representative of at least 3 independent experiments.

**Fig. S4.**
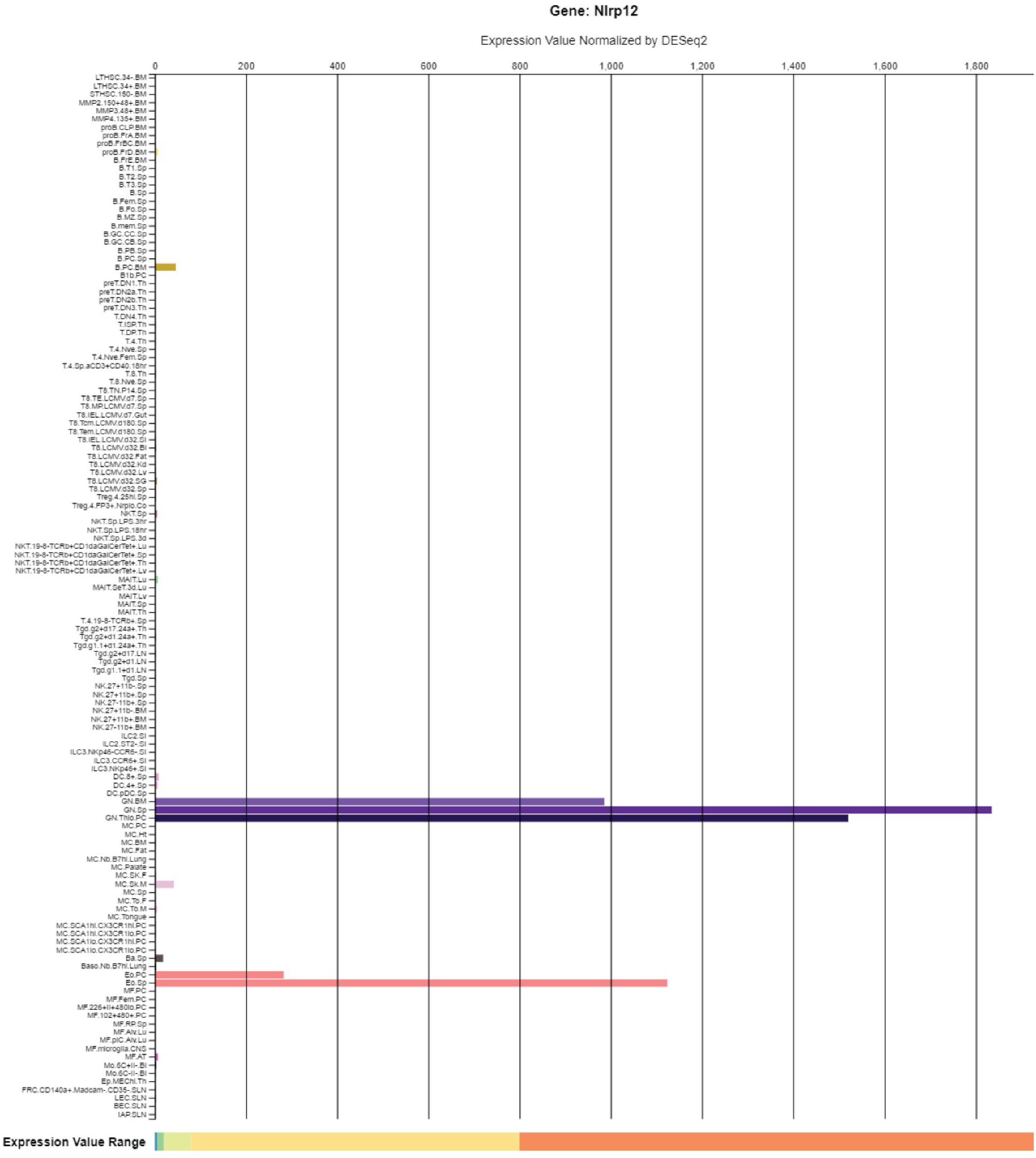
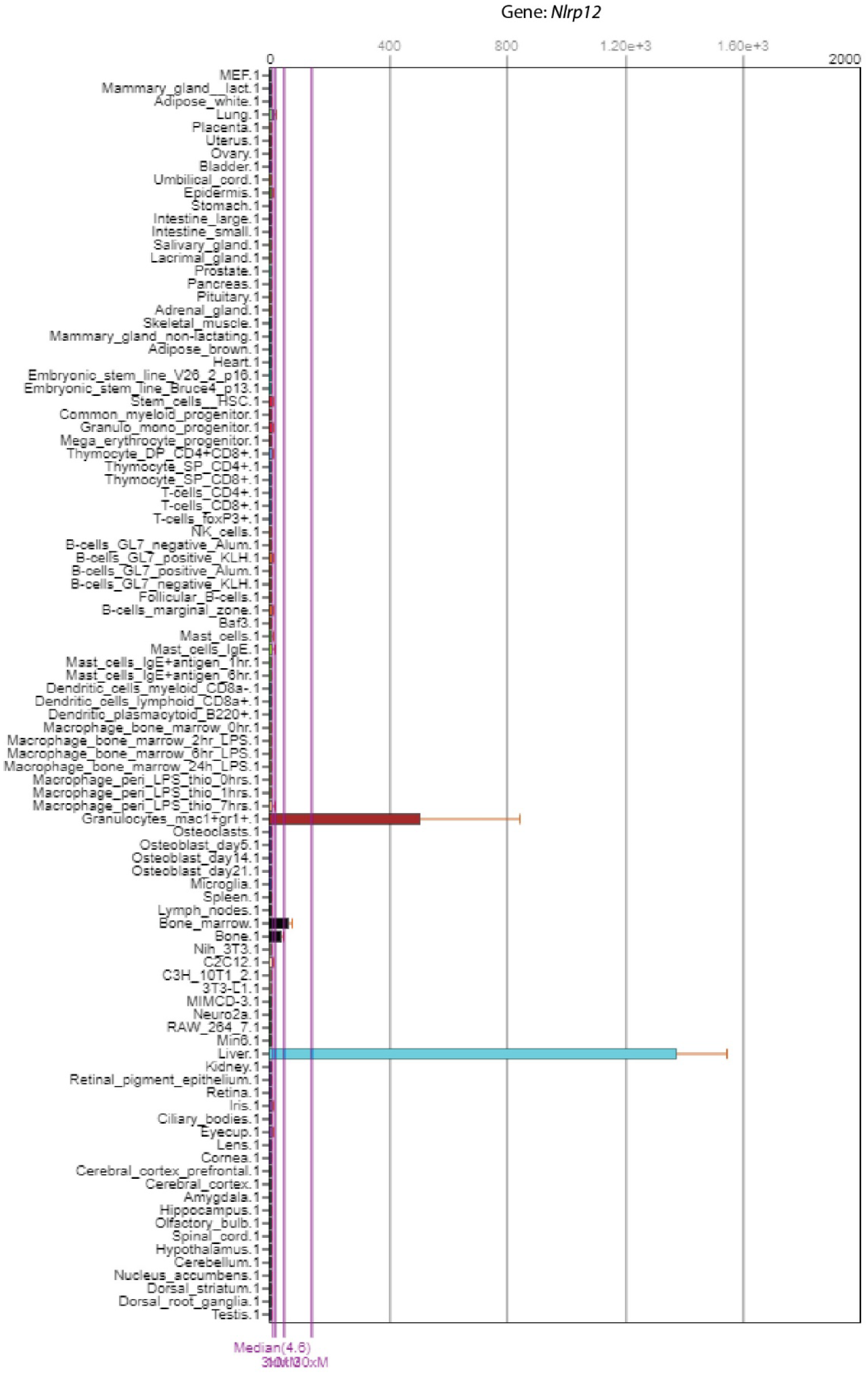
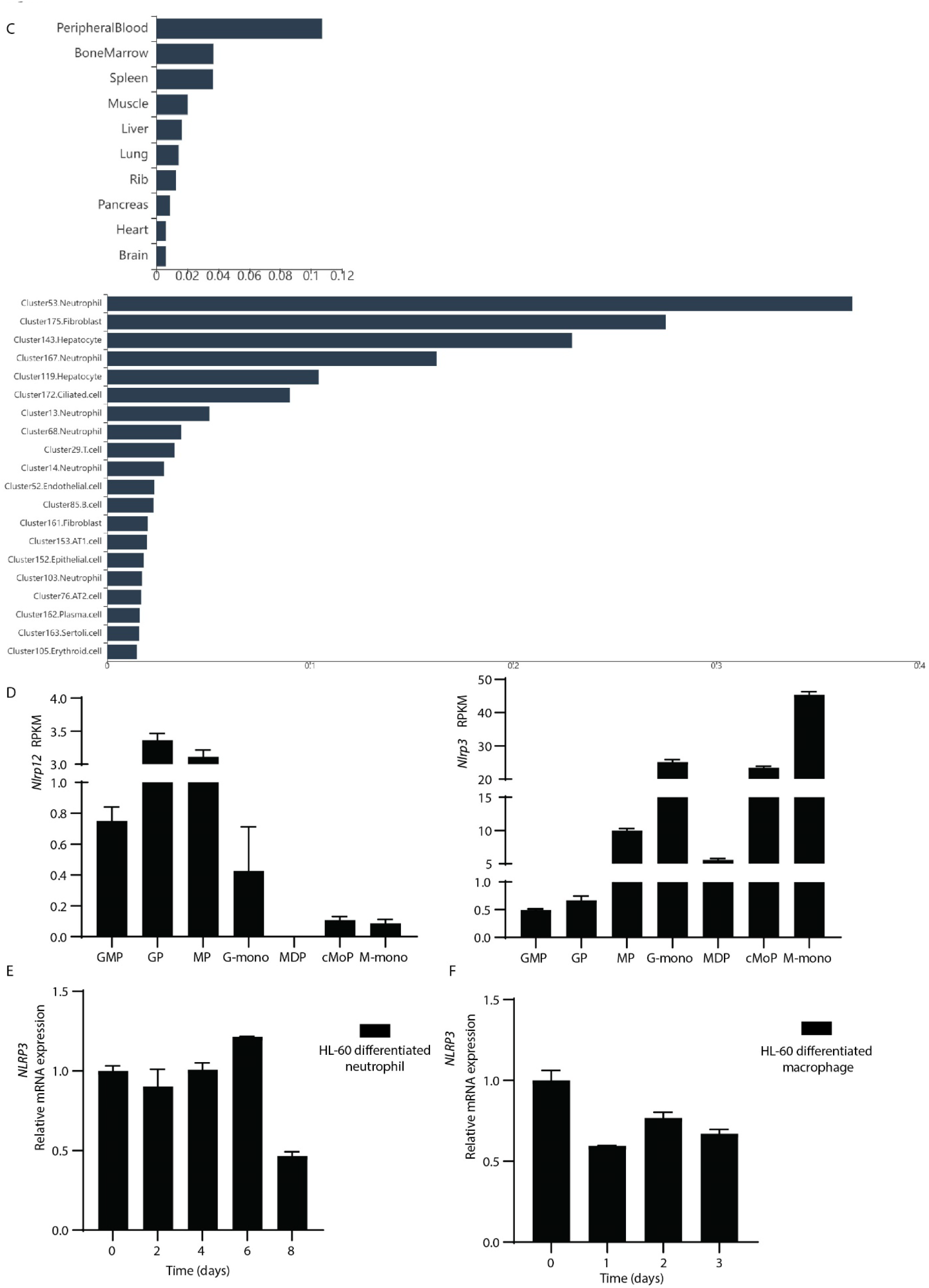
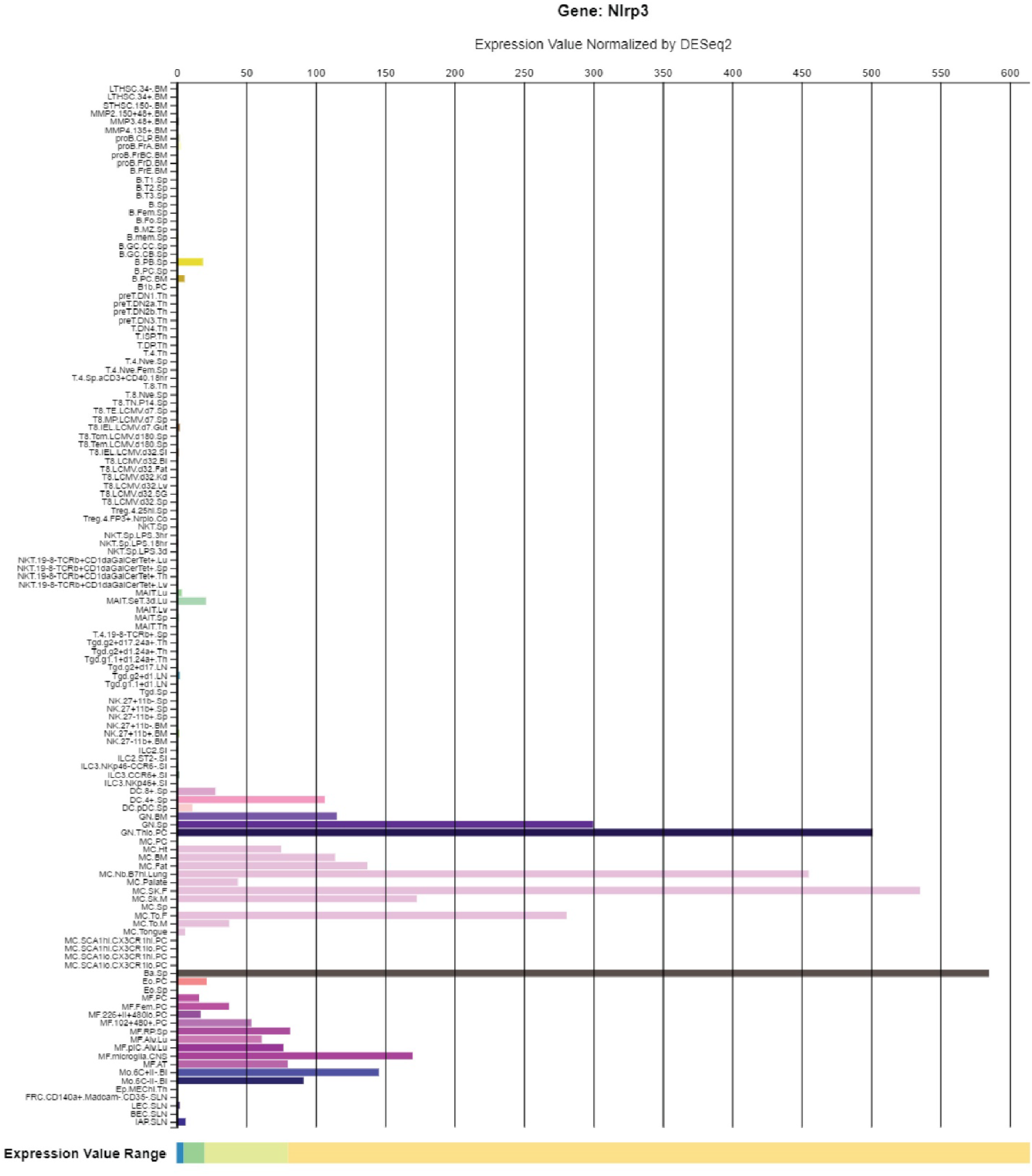
Specific expression of NLRP12 and NLRP3. (A-C) The expression of *Nlrp12* from ImmGen (A), BioGPS (B), and the Mouse Cell Altas (C). The results were downloaded from BioGPS, ImmGen (RNA-seq) and the Mouse Cell Altas websites. (D) The expression of *Nlrp3* and *Nlrp12* from Fig. 4B depicted as a bar graph. (E, F) qRT-PCR analysis of *NLRP3* expression. The samples in Fig. 4E (E) and Fig. 4F (F) were assayed for *NLRP3* expression. (G) The expression of *Nlrp3* from ImmGen. Data from lung basophils from mice infected with *Nippostrongylus brasiliensis* were excluded from the graph as these had extraordinarily high expression of *Nlrp3* that obscured visualization of the basal expression in other cell types. The results in (D), (E) and (F) are representative of at least 3 independent experiments.

**Fig. S5.**
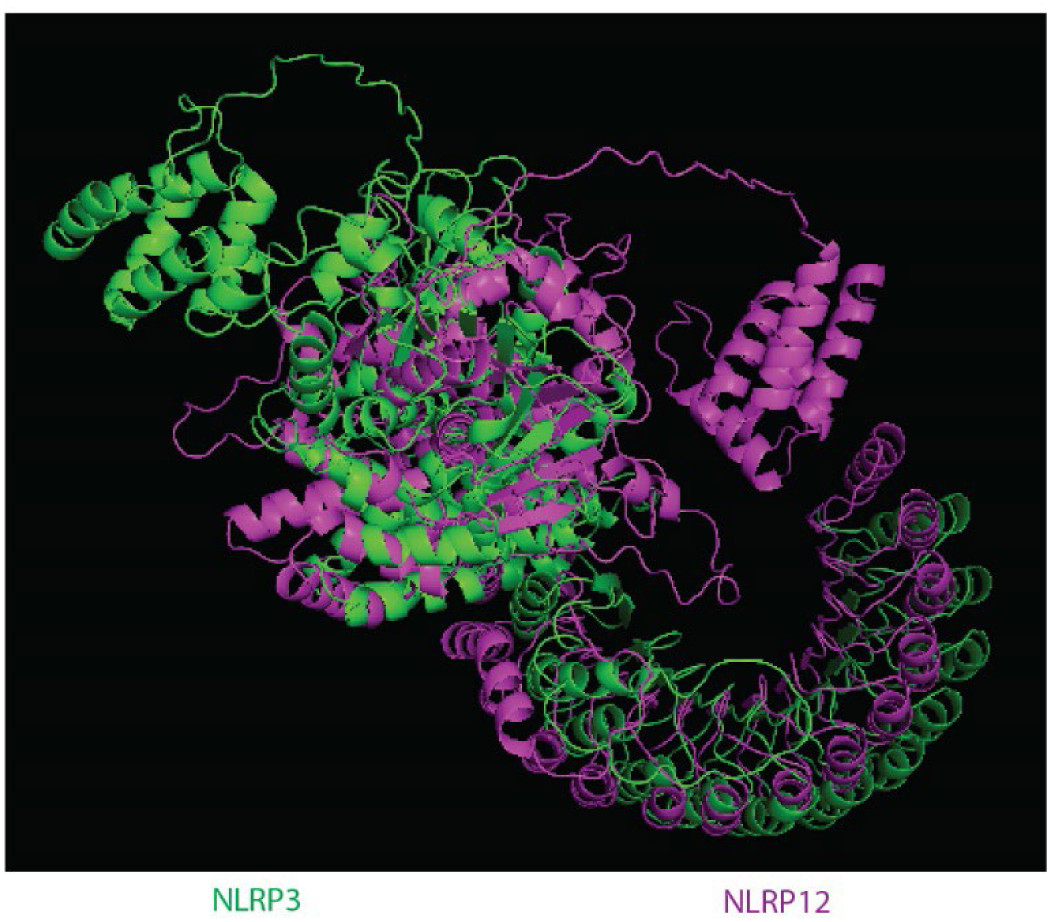
Alignment the NLRP12 structure with NLRP3 structure. (A) The structure files of mouse NLRP3 (green) and NLRP12 (magenta) were downloaded from AlphaFold protein structure database and aligned by PyMOL.

**Fig. S6.**
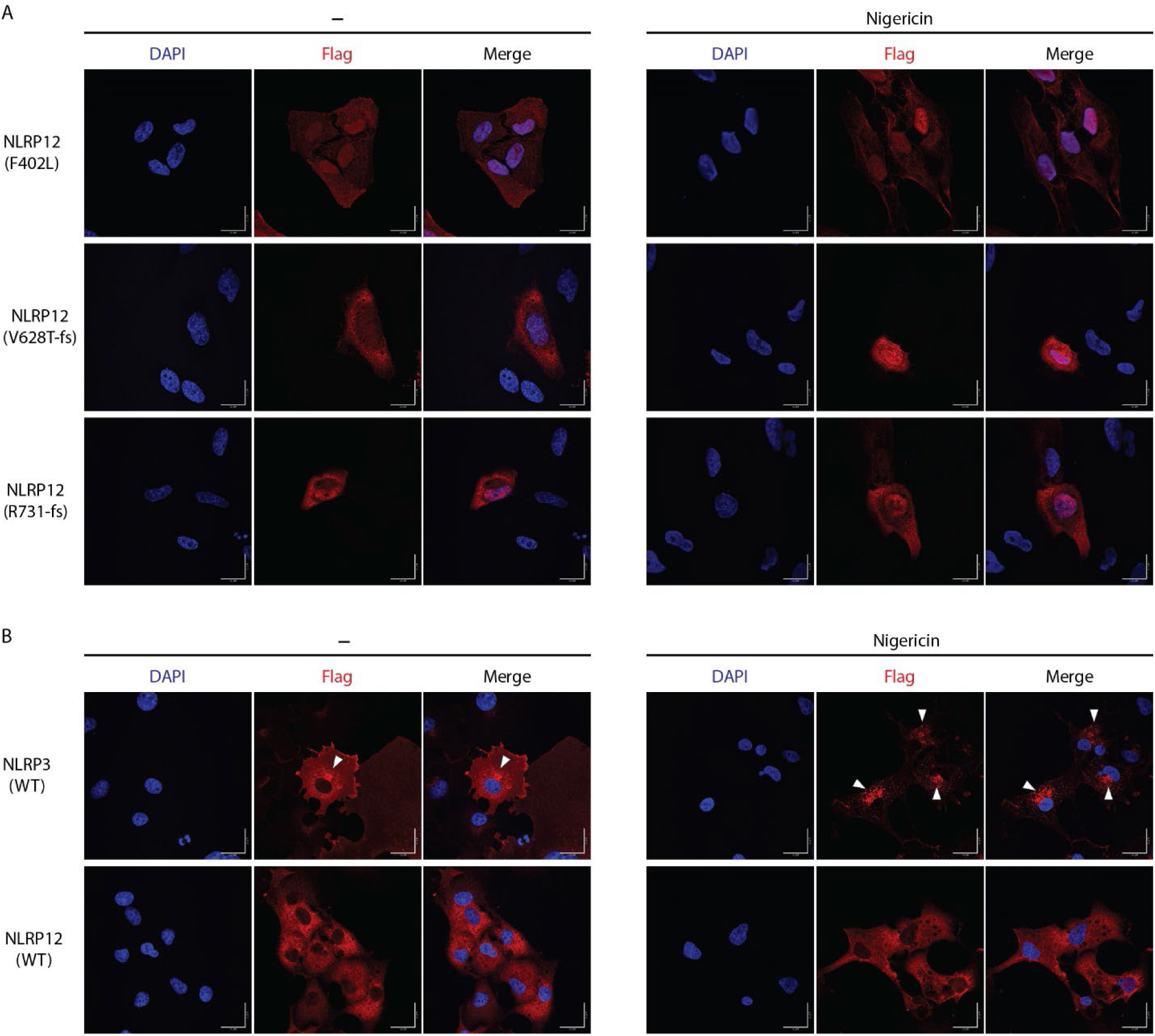
The subcellular location of NLRP3 and NLRP12. (A) The subcellular location of NLRP12(F402L, V628T-fs, R731-fs) in HeLa stable cells. The HeLa cells stably expressing NLRP12 were stimulated with nigericin or not. Immunostaining was performed for the Flag epitope tag and cells visualized by confocal microscopy. Scale bar, 20μm. (B) The subcellular location of NLRP3 and NLRP12 in COS-1 stable cells. The COS-1 cells stably expressing mouse NLRP3 and NLRP12 were stimulated with nigericin or not. Immunostaining was performed for the Flag epitope tag and cells visualized by confocal microscopy. Scale bar, 20μm. Triangles indicate the NLRP3 foci. All the images are representative of at least 3 independent experiments.

**Table. S1.**
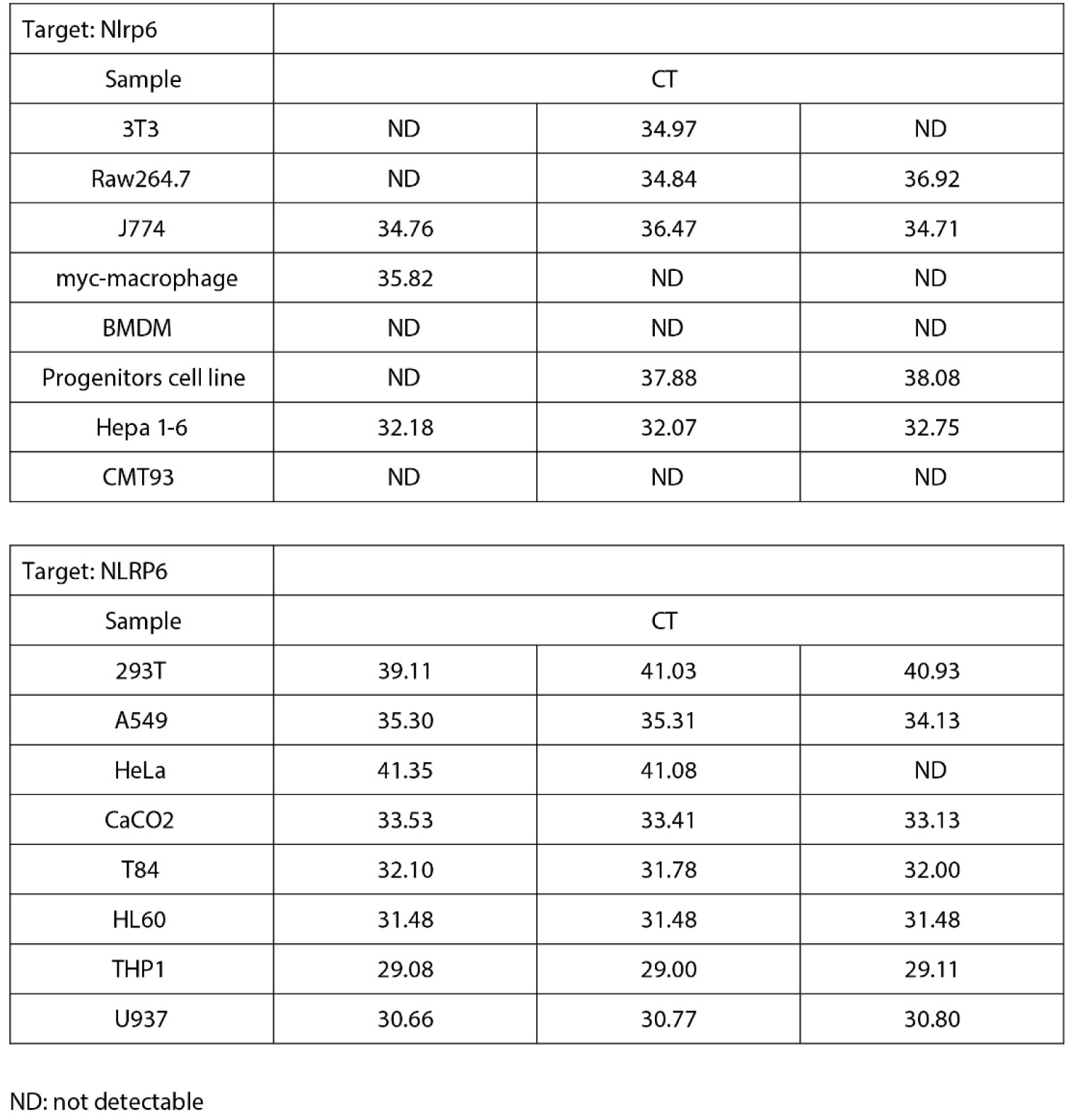
Expression of *Nlrp6* and *NLRP6* in various cell lines. qRT-PCR analysis of *Nlrp6* and *NLRP6* expression in indicated mouse and human cell lines. The table showed the raw CT value. All the results are representative of at least 3 independent experiments.

|  |  |  |  |
| --- | --- | --- | --- |
| Target: Nlrp6 |  |  |  |
| Sample | CT |  |  |
| 3T3 | ND | 34.97 | ND |
| Raw264.7 | ND | 34.84 | 36.92 |
| J774 | 34.76 | 36.47 | 34.71 |
| myc-macrophage | 35.82 | ND | ND |
| BMDM | ND | ND | ND |
| Progenitors cell line | ND | 37.88 | 38.08 |
| Hepa 1-6 | 32.18 | 32.07 | 32.75 |
| CMT93 | ND | ND | ND |
| Target: NLRP6 |  |  |  |
| Sample | CT |  |  |
| 293T | 39.11 | 41.03 | 40.93 |
| A549 | 35.30 | 35.31 | 34.13 |
| HeLa | 41.35 | 41.08 | ND |
| CaCO2 | 33.53 | 33.41 | 33.13 |
| T84 | 32.10 | 31.78 | 32.00 |
| HL60 | 31.48 | 31.48 | 31.48 |
| THP1 | 29.08 | 29.00 | 29.11 |
| U937 | 30.66 | 30.77 | 30.80 |

ND: not detectable

## Reference and Notes

1. S. B. Kovacs, E. A. Miao, Gasdermins: Effectors of Pyroptosis. Trends Cell Biol 27, 673–684 (2017).

2. K. Nozaki, L. Li, E. A. Miao, Innate Sensors Trigger Regulated Cell Death to Combat Intracellular Infection. Annu Rev Immunol 40, 469–498 (2022).

3. C. A. Janeway, Jr., Approaching the asymptote? Evolution and revolution in immunology. Cold Spring Harb Symp Quant Biol 54 **Pt** **1**, 1–13 (1989).

4. P. Devant, J. C. Kagan, Molecular mechanisms of gasdermin D pore-forming activity. Nat Immunol 24, 1064–1075 (2023).

5. S. Xia, Z. Zhang, V. G. Magupalli, J. L. Pablo, Y. Dong, S. M. Vora, L. Wang, T. M. Fu, M. P. Jacobson, A. Greka, J. Lieberman, J. Ruan, H. Wu, Gasdermin D pore structure reveals preferential release of mature interleukin-1. Nature 593, 607–611 (2021).

6. J. A. Hagar, D. A. Powell, Y. Aachoui, R. K. Ernst, E. A. Miao, Cytoplasmic LPS activates caspase-11: implications in TLR4-independent endotoxic shock. Science 341, 1250–1253 (2013).

7. J. Shi, Y. Zhao, Y. Wang, W. Gao, J. Ding, P. Li, L. Hu, F. Shao, Inflammatory caspases are innate immune receptors for intracellular LPS. Nature 514, 187–192 (2014).

8. N. Kayagaki, M. T. Wong, I. B. Stowe, S. R. Ramani, L. C. Gonzalez, S. Akashi-Takamura, K. Miyake, J. Zhang, W. P. Lee, A. Muszynski, L. S. Forsberg, R. W. Carlson, V. M. Dixit, Noncanonical inflammasome activation by intracellular LPS independent of TLR4. Science 341, 1246–1249 (2013).

9. J. A. Duncan, S. W. Canna, The NLRC4 Inflammasome. Immunol Rev 281, 115–123 (2018).

10. K. V. Swanson, M. Deng, J. P. Ting, The NLRP3 inflammasome: molecular activation and regulation to therapeutics. Nat Rev Immunol 19, 477–489 (2019).

11. Y. He, M. Y. Zeng, D. Yang, B. Motro, G. Nunez, NEK7 is an essential mediator of NLRP3 activation downstream of potassium efflux. Nature 530, 354–357 (2016).

12. H. Shi, Y. Wang, X. Li, X. Zhan, M. Tang, M. Fina, L. Su, D. Pratt, C. H. Bu, S. Hildebrand, S. Lyon, L. Scott, J. Quan, Q. Sun, J. Russell, S. Arnett, P. Jurek, D. Chen, V. V. Kravchenko, J. C. Mathison, E. M. Moresco, N. L. Monson, R. J. Ulevitch, B. Beutler, NLRP3 activation and mitosis are mutually exclusive events coordinated by NEK7, a new inflammasome component. Nat Immunol 17, 250–258 (2016).

13. J. L. Schmid-Burgk, D. Chauhan, T. Schmidt, T. S. Ebert, J. Reinhardt, E. Endl, V. Hornung, A Genome-wide CRISPR (Clustered Regularly Interspaced Short Palindromic Repeats) Screen Identifies NEK7 as an Essential Component of NLRP3 Inflammasome Activation. J Biol Chem 291, 103–109 (2016).

14. J. P. Ting, R. C. Lovering, E. S. Alnemri, J. Bertin, J. M. Boss, B. K. Davis, R. A. Flavell, S. E. Girardin, A. Godzik, J. A. Harton, H. M. Hoffman, J. P. Hugot, N. Inohara, A. Mackenzie, L. J. Maltais, G. Nunez, Y. Ogura, L. A. Otten, D. Philpott, J. C. Reed, W. Reith, S. Schreiber, V. Steimle, P. A. Ward, The NLR gene family: a standard nomenclature. Immunity 28, 285–287 (2008).

15. H. H. Park, Y. C. Lo, S. C. Lin, L. Wang, J. K. Yang, H. Wu, The death domain superfamily in intracellular signaling of apoptosis and inflammation. Annu Rev Immunol 25, 561–586 (2007).

16. J. Masumoto, T. A. Dowds, P. Schaner, F. F. Chen, Y. Ogura, M. Li, L. Zhu, T. Katsuyama, J. Sagara, S. Taniguchi, D. L. Gumucio, G. Nunez, N. Inohara, ASC is an activating adaptor for NF-kappa B and caspase-8-dependent apoptosis. Biochem Biophys Res Commun 303, 69–73 (2003).

17. S. Zhu, S. Ding, P. Wang, Z. Wei, W. Pan, N. W. Palm, Y. Yang, H. Yu, H. B. Li, G. Wang, X. Lei, M. R. de Zoete, J. Zhao, Y. Zheng, H. Chen, Y. Zhao, K. A. Jurado, N. Feng, L. Shan, Y. Kluger, J. Lu, C. Abraham, E. Fikrig, H. B. Greenberg, R. A. Flavell, Nlrp9b inflammasome restricts rotavirus infection in intestinal epithelial cells. Nature 546, 667–670 (2017).

18. D. Zheng, G. Mohapatra, L. Kern, Y. He, M. D. Shmueli, R. Valdes-Mas, A. A. Kolodziejczyk, T. Prochnicki, M. B. Vasconcelos, L. Schorr, F. Hertel, Y. S. Lee, M. C. Rufino, E. Ceddaha, S. Shimshy, R. J. Hodgetts, M. Dori-Bachash, C. Kleimeyer, K. Goldenberg, M. Heinemann, N. Stettner, A. Harmelin, H. Shapiro, J. Puschhof, M. Chen, R. A. Flavell, E. Latz, Y. Merbl, S. K. Abdeen, E. Elinav, Epithelial Nlrp10 inflammasome mediates protection against intestinal autoinflammation. Nat Immunol 24, 585–594 (2023).

19. T. Prochnicki, M. B. Vasconcelos, K. S. Robinson, M. S. J. Mangan, D. De Graaf, K. Shkarina, M. Lovotti, L. Standke, R. Kaiser, R. Stahl, F. G. Duthie, M. Rothe, K. Antonova, L. M. Jenster, Z. H. Lau, S. Rosing, N. Mirza, C. Gottschild, D. Wachten, C. Gunther, T. A. Kufer, F. I. Schmidt, F. L. Zhong, E. Latz, Mitochondrial damage activates the NLRP10 inflammasome. Nat Immunol 24, 595–603 (2023).

20. S. Khare, A. Dorfleutner, N. B. Bryan, C. Yun, A. D. Radian, L. de Almeida, Y. Rojanasakul, C. Stehlik, An NLRP7-containing inflammasome mediates recognition of microbial lipopeptides in human macrophages. Immunity 36, 464–476 (2012).

21. A. Akbal, A. Dernst, M. Lovotti, M. S. J. Mangan, R. M. McManus, E. Latz, How location and cellular signaling combine to activate the NLRP3 inflammasome. Cell Mol Immunol 19, 1201–1214 (2022).

22. S. Zangiabadi, A. A. Abdul-Sater, Regulation of the NLRP3 Inflammasome by Posttranslational Modifications. J Immunol 208, 286–292 (2022).

23. H. Shi, A. Murray, B. Beutler, Reconstruction of the Mouse Inflammasome System in HEK293T Cells. Bio Protoc 6, (2016).

24. G. Meng, F. Zhang, I. Fuss, A. Kitani, W. Strober, A mutation in the Nlrp3 gene causing inflammasome hyperactivation potentiates Th17 cell-dominant immune responses. Immunity 30, 860–874 (2009).

25. H. Hara, S. S. Seregin, D. Yang, K. Fukase, M. Chamaillard, E. S. Alnemri, N. Inohara, G. Y. Chen, G. Nunez, The NLRP6 Inflammasome Recognizes Lipoteichoic Acid and Regulates Gram-Positive Pathogen Infection. Cell 175, 1651–1664 e1614 (2018).

26. D. H. Reikvam, A. Erofeev, A. Sandvik, V. Grcic, F. L. Jahnsen, P. Gaustad, K. D. McCoy, A. J. Macpherson, L. A. Meza-Zepeda, F. E. Johansen, Depletion of murine intestinal microbiota: effects on gut mucosa and epithelial gene expression. PLoS One 6, e17996 (2011).

27. S. Borghini, S. Tassi, S. Chiesa, F. Caroli, S. Carta, R. Caorsi, M. Fiore, L. Delfino, D. Lasiglie, C. Ferraris, E. Traggiai, M. Di Duca, G. Santamaria, A. D’Osualdo, M. Tosca, A. Martini, I. Ceccherini, A. Rubartelli, M. Gattorno, Clinical presentation and pathogenesis of cold-induced autoinflammatory disease in a family with recurrence of an NLRP12 mutation. Arthritis Rheum 63, 830–839 (2011).

28. C. De Pieri, J. Vuch, E. Athanasakis, G. M. Severini, S. Crovella, A. M. Bianco, A. Tommasini, F402L variant in NLRP12 in subjects with undiagnosed periodic fevers and in healthy controls. Clin Exp Rheumatol 32, 993–994 (2014).

29. N. Jacob, S. S. Dasharathy, V. Bui, J. N. Benhammou, W. W. Grody, R. R. Singh, J. R. Pisegna, Generalized Cytokine Increase in the Setting of a Multisystem Clinical Disorder and Carcinoid Syndrome Associated with a Novel NLRP12 Variant. Dig Dis Sci 64, 2140–2146 (2019).

30. I. Jeru, G. Le Borgne, E. Cochet, H. Hayrapetyan, P. Duquesnoy, G. Grateau, A. Morali, T. Sarkisian, S. Amselem, Identification and functional consequences of a recurrent NLRP12 missense mutation in periodic fever syndromes. Arthritis Rheum 63, 1459–1464 (2011).

31. I. Jeru, P. Duquesnoy, T. Fernandes-Alnemri, E. Cochet, J. W. Yu, M. Lackmy-Port-Lis, E. Grimprel, J. Landman-Parker, V. Hentgen, S. Marlin, K. McElreavey, T. Sarkisian, G. Grateau, E. S. Alnemri, S. Amselem, Mutations in NALP12 cause hereditary periodic fever syndromes. Proc Natl Acad Sci U S A 105, 1614–1619 (2008).

32. M. Shen, L. Tang, X. Shi, X. Zeng, Q. Yao, NLRP12 autoinflammatory disease: a Chinese case series and literature review. Clin Rheumatol 36, 1661–1667 (2017).

33. S. Borte, M. H. Celiksoy, V. Menzel, O. Ozkaya, F. Z. Ozen, L. Hammarstrom, A. Yildiran, Novel NLRP12 mutations associated with intestinal amyloidosis in a patient diagnosed with common variable immunodeficiency. Clin Immunol 154, 105–111 (2014).

34. A. Vitale, D. Rigante, M. C. Maggio, G. Emmi, M. Romano, E. Silvestri, O. M. Lucherini, L. Emmi, V. Gerloni, L. Cantarini, Rare NLRP12 variants associated with the NLRP12-autoinflammatory disorder phenotype: an Italian case series. Clin Exp Rheumatol 31, 155–156 (2013).

35. A. Vitale, D. Rigante, O. M. Lucherini, F. Caso, L. Cantarini, The role of the F402L allele in the NLRP12-autoinflammatory disorder. Reply to: F402L variant in NLRP12 in subjects with undiagnosed periodic fevers and in healthy controls, De Pieri et al. Clin Exp Rheumatol 32, 994 (2014).

36. M. M. Kostik, E. N. Suspitsin, M. N. Guseva, A. S. Levina, A. Y. Kazantseva, A. P. Sokolenko, E. N. Imyanitov, Multigene sequencing reveals heterogeneity of NLRP12-related autoinflammatory disorders. Rheumatol Int 38, 887–893 (2018).

37. W. Wang, Y. Zhou, L. Q. Zhong, Z. Li, S. Jian, X. Y. Tang, H. M. Song, The clinical phenotype and genotype of NLRP12-autoinflammatory disease: a Chinese case series with literature review. World J Pediatr 16, 514–519 (2020).

38. G. G. Wang, K. R. Calvo, M. P. Pasillas, D. B. Sykes, H. Hacker, M. P. Kamps, Quantitative production of macrophages or neutrophils ex vivo using conditional Hoxb8. Nat Methods 3, 287–293 (2006).

39. R. A. Fleck, S. Romero-Steiner, M. H. Nahm, Use of HL-60 cell line to measure opsonic capacity of pneumococcal antibodies. Clin Diagn Lab Immunol 12, 19–27 (2005).

40. D. Gupta, H. P. Shah, K. Malu, N. Berliner, P. Gaines, Differentiation and characterization of myeloid cells. Curr Protoc Immunol 104, 22F 25 21–22F 25 28 (2014).

41. J. Chen, Z. J. Chen, PtdIns4P on dispersed trans-Golgi network mediates NLRP3 inflammasome activation. Nature 564, 71–76 (2018).

42. Z. Zhang, R. Venditti, L. Ran, Z. Liu, K. Vivot, A. Schurmann, J. S. Bonifacino, M. A. De Matteis, R. Ricci, Distinct changes in endosomal composition promote NLRP3 inflammasome activation. Nat Immunol 24, 30–41 (2023).

43. L. W. Peterson, D. Artis, Intestinal epithelial cells: regulators of barrier function and immune homeostasis. Nat Rev Immunol 14, 141–153 (2014).

44. R. Okumura, K. Takeda, Roles of intestinal epithelial cells in the maintenance of gut homeostasis. Exp Mol Med 49, e338 (2017).

45. C. Sauvanet, J. Wayt, T. Pelaseyed, A. Bretscher, Structure, regulation, and functional diversity of microvilli on the apical domain of epithelial cells. Annu Rev Cell Dev Biol 31, 593–621 (2015).

46. J. D. Jones, R. E. Vance, J. L. Dangl, Intracellular innate immune surveillance devices in plants and animals. Science 354, (2016).

47. W. M. Nauseef, R. A. Clark, in Mandell, Douglas, and Bennett’s Principles and Practice of Infectious Diseases. (2010), pp. 99–127.

48. J. D. Lich, K. L. Williams, C. B. Moore, J. C. Arthur, B. K. Davis, D. J. Taxman, J. P. Ting, Monarch-1 suppresses non-canonical NF-kappaB activation and p52-dependent chemokine expression in monocytes. J Immunol 178, 1256–1260 (2007).

49. I. C. Allen, J. E. Wilson, M. Schneider, J. D. Lich, R. A. Roberts, J. C. Arthur, R. M. Woodford, B. K. Davis, J. M. Uronis, H. H. Herfarth, C. Jobin, A. B. Rogers, J. P. Ting, NLRP12 suppresses colon inflammation and tumorigenesis through the negative regulation of noncanonical NF-kappaB signaling. Immunity 36, 742–754 (2012).

50. M. H. Zaki, P. Vogel, R. K. Malireddi, M. Body-Malapel, P. K. Anand, J. Bertin, D. R. Green, M. Lamkanfi, T. D. Kanneganti, The NOD-like receptor NLRP12 attenuates colon inflammation and tumorigenesis. Cancer Cell 20, 649–660 (2011).

51. S. N. Udden, Y. T. Kwak, V. Godfrey, M. A. W. Khan, S. Khan, N. Loof, L. Peng, H. Zhu, H. Zaki, NLRP12 suppresses hepatocellular carcinoma via downregulation of cJun N-terminal kinase activation in the hepatocyte. Elife 8, (2019).

52. T. K. Ulland, N. Jain, E. E. Hornick, E. I. Elliott, G. M. Clay, J. J. Sadler, K. A. Mills, A. M. Janowski, A. P. Volk, K. Wang, K. L. Legge, L. Gakhar, M. Bourdi, P. J. Ferguson, M. E. Wilson, S. L. Cassel, F. S. Sutterwala, Nlrp12 mutation causes C57BL/6J strain-specific defect in neutrophil recruitment. Nat Commun 7, 13180 (2016).

53. G. I. Vladimer, D. Weng, S. W. Paquette, S. K. Vanaja, V. A. Rathinam, M. H. Aune, J. E. Conlon, J. J. Burbage, M. K. Proulx, Q. Liu, G. Reed, J. C. Mecsas, Y. Iwakura, J. Bertin, J. D. Goguen, K. A. Fitzgerald, E. Lien, The NLRP12 inflammasome recognizes Yersinia pestis. Immunity 37, 96–107 (2012).

54. J. R. Coombs, A. Zamoshnikova, C. L. Holley, M. P. Maddugoda, D. E. T. Teo, C. Chauvin, L. F. Poulin, N. Vitak, C. M. Ross, M. Mellacheruvu, R. C. Coll, L. X. Heinz, S. S. Burgener, S. Emming, M. Chamaillard, D. Boucher, K. Schroder, NLRP12 interacts with NLRP3 to block the activation of the human NLRP3 inflammasome. Sci Signal 17, eabg8145 (2024).

55. B. Sundaram, N. Pandian, R. Mall, Y. Wang, R. Sarkar, H. J. Kim, R. K. S. Malireddi, R. Karki, L. J. Janke, P. Vogel, T. D. Kanneganti, NLRP12-PANoptosome activates PANoptosis and pathology in response to heme and PAMPs. Cell 186, 2783–2801 e2720 (2023).

56. M. Centola, G. Wood, D. M. Frucht, J. Galon, M. Aringer, C. Farrell, D. W. Kingma, M. E. Horwitz, E. Mansfield, S. M. Holland, J. J. O’Shea, H. F. Rosenberg, H. L. Malech, D. L. Kastner, The gene for familial Mediterranean fever, MEFV, is expressed in early leukocyte development and is regulated in response to inflammatory mediators. Blood 95, 3223–3231 (2000).

57. S. Seshadri, M. D. Duncan, J. M. Hart, M. A. Gavrilin, M. D. Wewers, Pyrin levels in human monocytes and monocyte-derived macrophages regulate IL-1beta processing and release. J Immunol 179, 1274–1281 (2007).

58. I. Jeru, V. Hentgen, S. Normand, P. Duquesnoy, E. Cochet, A. Delwail, G. Grateau, S. Marlin, S. Amselem, J. C. Lecron, Role of interleukin-1beta in NLRP12-associated autoinflammatory disorders and resistance to anti-interleukin-1 therapy. Arthritis Rheum 63, 2142–2148 (2011).

59. D. Novick, C. A. Dinarello, IL-18 binding protein reverses the life-threatening hyperinflammation of a baby with the NLRC4 mutation. J Allergy Clin Immunol 140, 316 (2017).

60. J. J. Hu, X. Liu, S. Xia, Z. Zhang, Y. Zhang, J. Zhao, J. Ruan, X. Luo, X. Lou, Y. Bai, J. Wang, L. R. Hollingsworth, V. G. Magupalli, L. Zhao, H. R. Luo, J. Kim, J. Lieberman, H. Wu, FDA-approved disulfiram inhibits pyroptosis by blocking gasdermin D pore formation. Nat Immunol, (2020).

61. J. Jumper, R. Evans, A. Pritzel, T. Green, M. Figurnov, O. Ronneberger, K. Tunyasuvunakool, R. Bates, A. Zidek, A. Potapenko, A. Bridgland, C. Meyer, S. A. A. Kohl, A. J. Ballard, A. Cowie, B. Romera-Paredes, S. Nikolov, R. Jain, J. Adler, T. Back, S. Petersen, D. Reiman, E. Clancy, M. Zielinski, M. Steinegger, M. Pacholska, T. Berghammer, S. Bodenstein, D. Silver, O. Vinyals, A. W. Senior, K. Kavukcuoglu, P. Kohli, D. Hassabis, Highly accurate protein structure prediction with AlphaFold. Nature 596, 583–589 (2021).

62. M. Varadi, S. Anyango, M. Deshpande, S. Nair, C. Natassia, G. Yordanova, D. Yuan, O. Stroe, G. Wood, A. Laydon, A. Zidek, T. Green, K. Tunyasuvunakool, S. Petersen, J. Jumper, E. Clancy, R. Green, A. Vora, M. Lutfi, M. Figurnov, A. Cowie, N. Hobbs, P. Kohli, G. Kleywegt, E. Birney, D. Hassabis, S. Velankar, AlphaFold Protein Structure Database: massively expanding the structural coverage of protein-sequence space with high-accuracy models. Nucleic Acids Res 50, D439–D444 (2022).

63. A. Yanez, S. G. Coetzee, A. Olsson, D. E. Muench, B. P. Berman, D. J. Hazelett, N. Salomonis, H. L. Grimes, H. S. Goodridge, Granulocyte-Monocyte Progenitors and Monocyte-Dendritic Cell Progenitors Independently Produce Functionally Distinct Monocytes. Immunity 47, 890–902 e894 (2017).

