## Supplementary figures and images for "NLRP3, NLRP6, and NLRP12 are inflammasomes with distinct expression patterns"

### Supplemental Figure 1

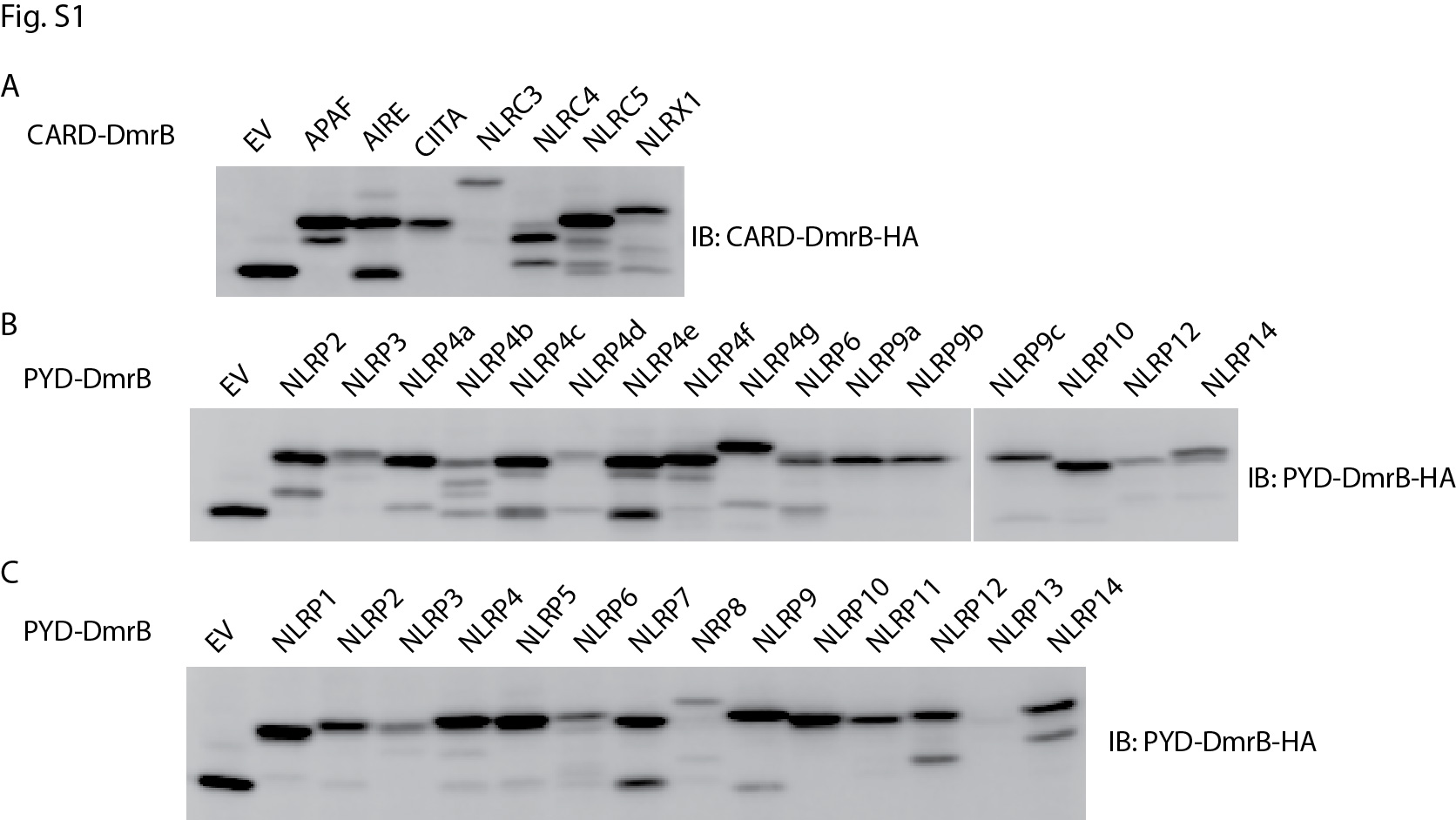

### Supplemental Figure 2

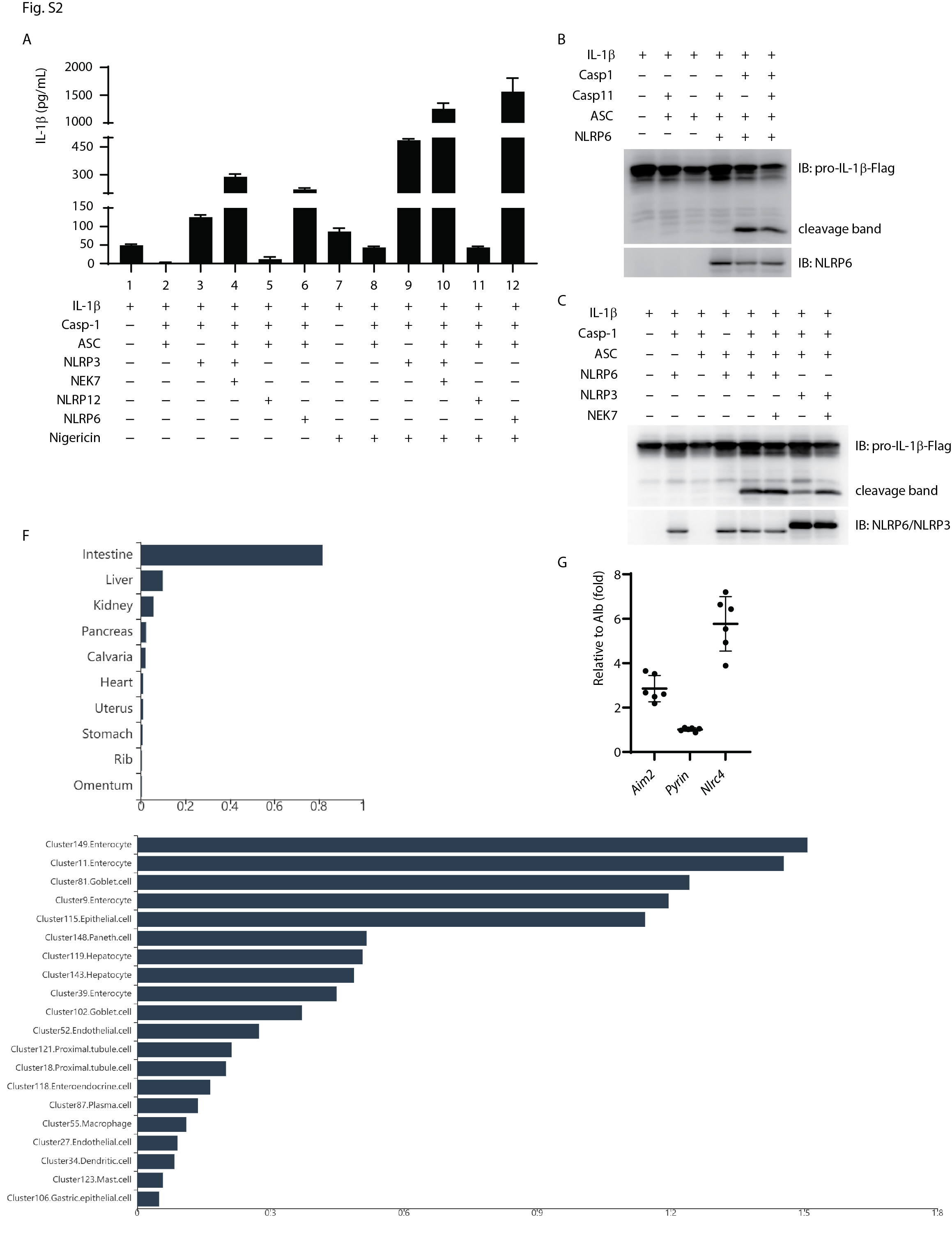

### Supplemental Figure 2D

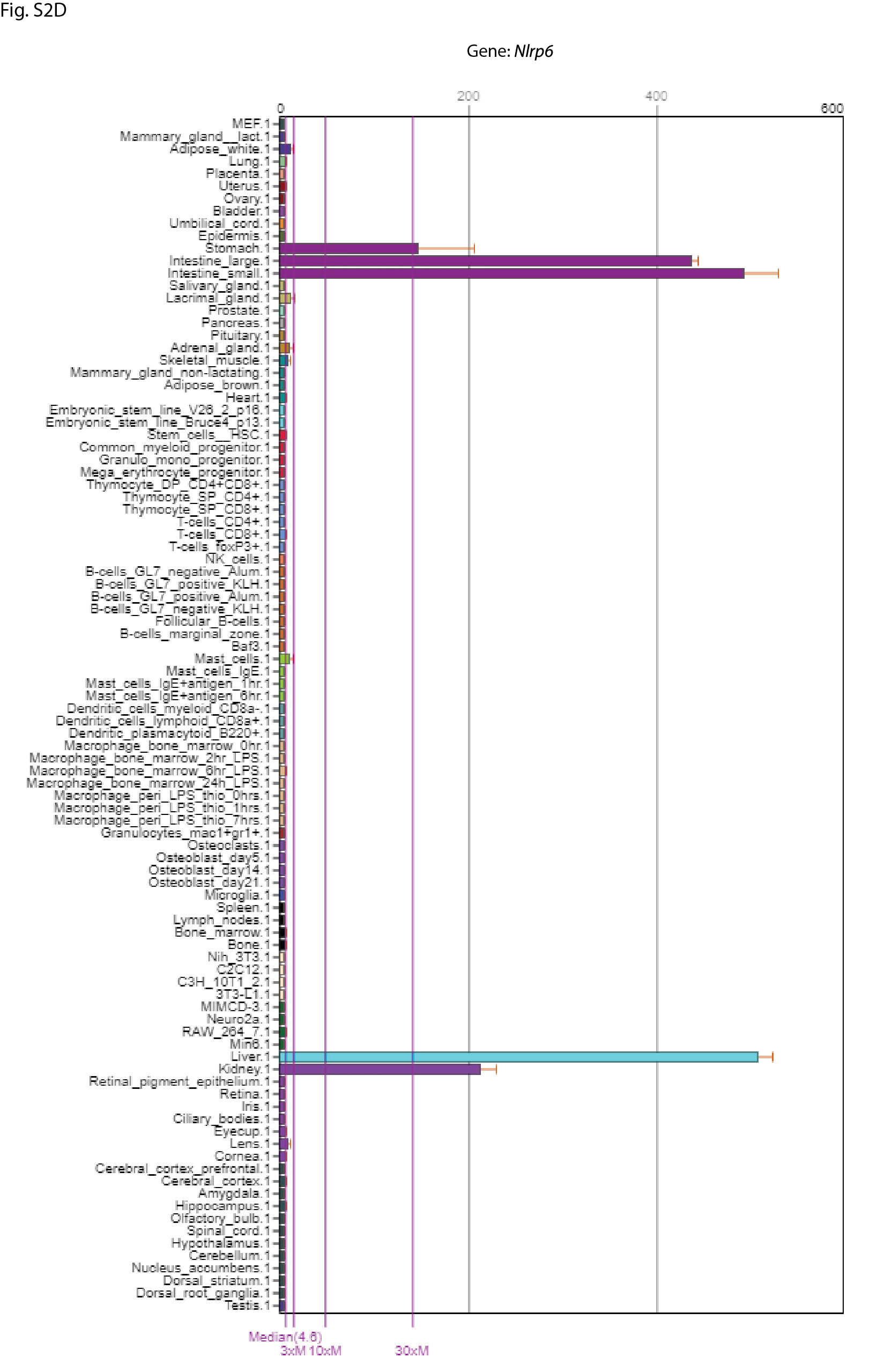

### Supplemental Figure 2E

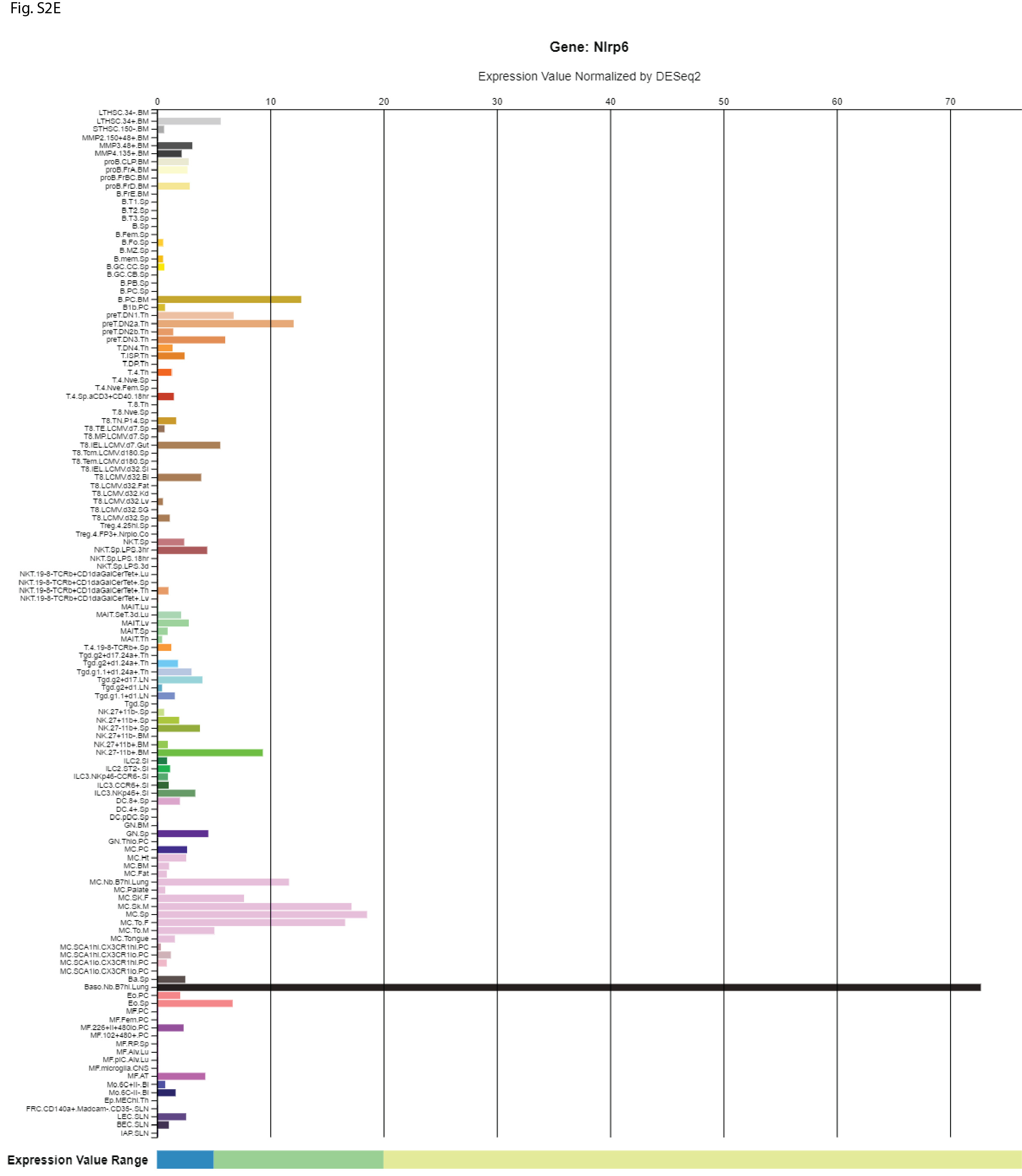

### Supplemental Figure 3

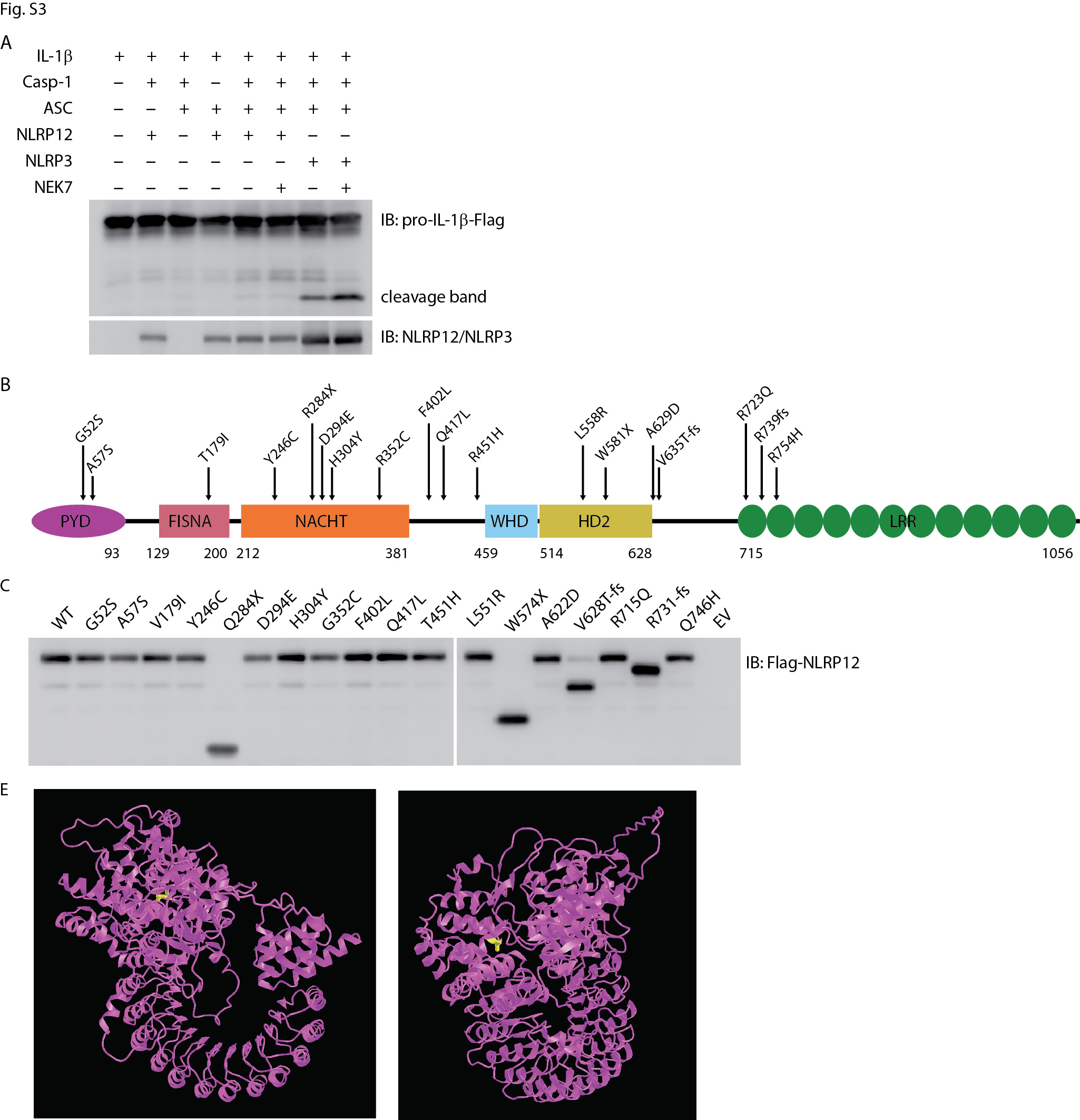

### Supplemental Figure 3D

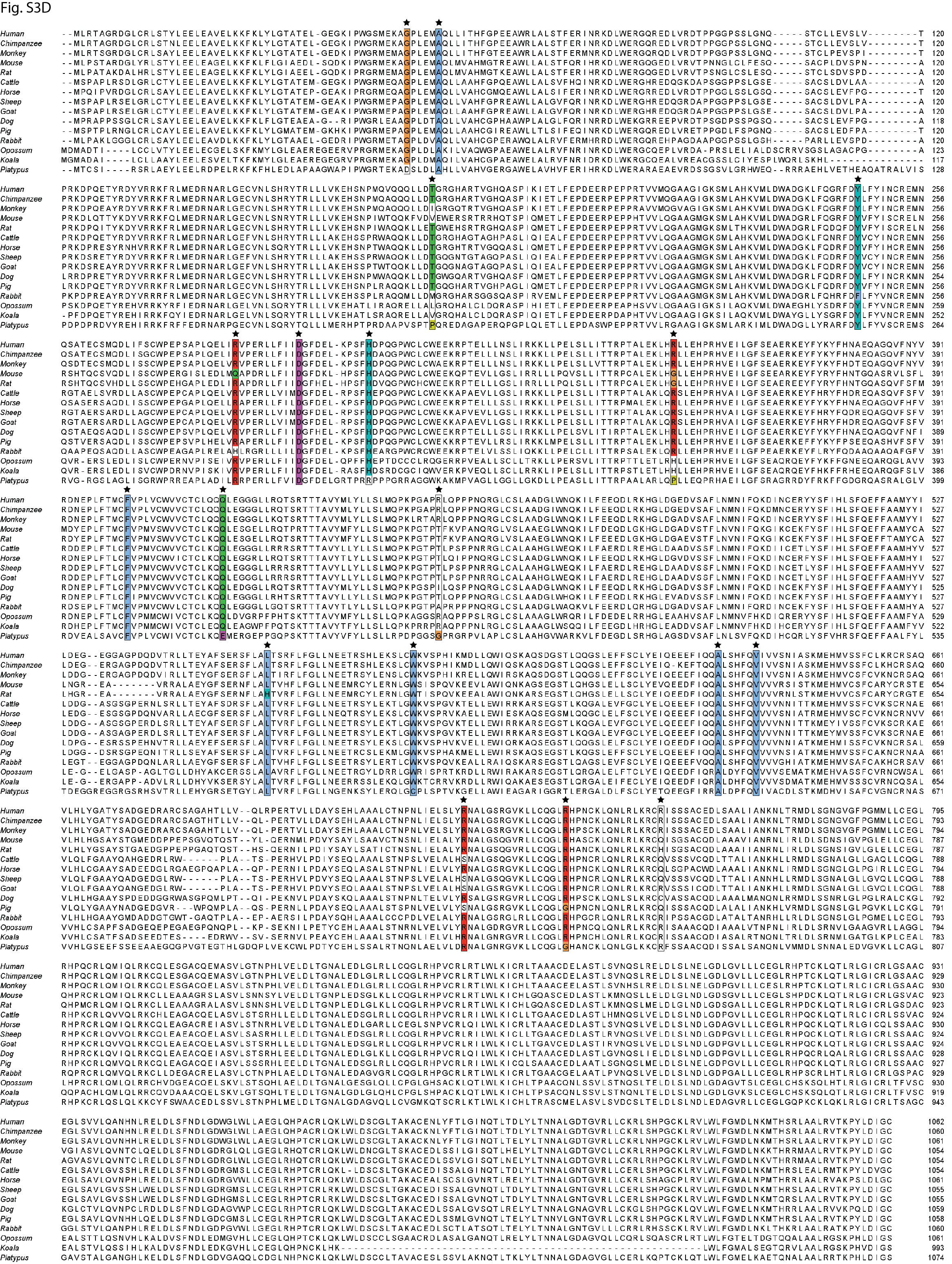

### Supplemental Figure 4A

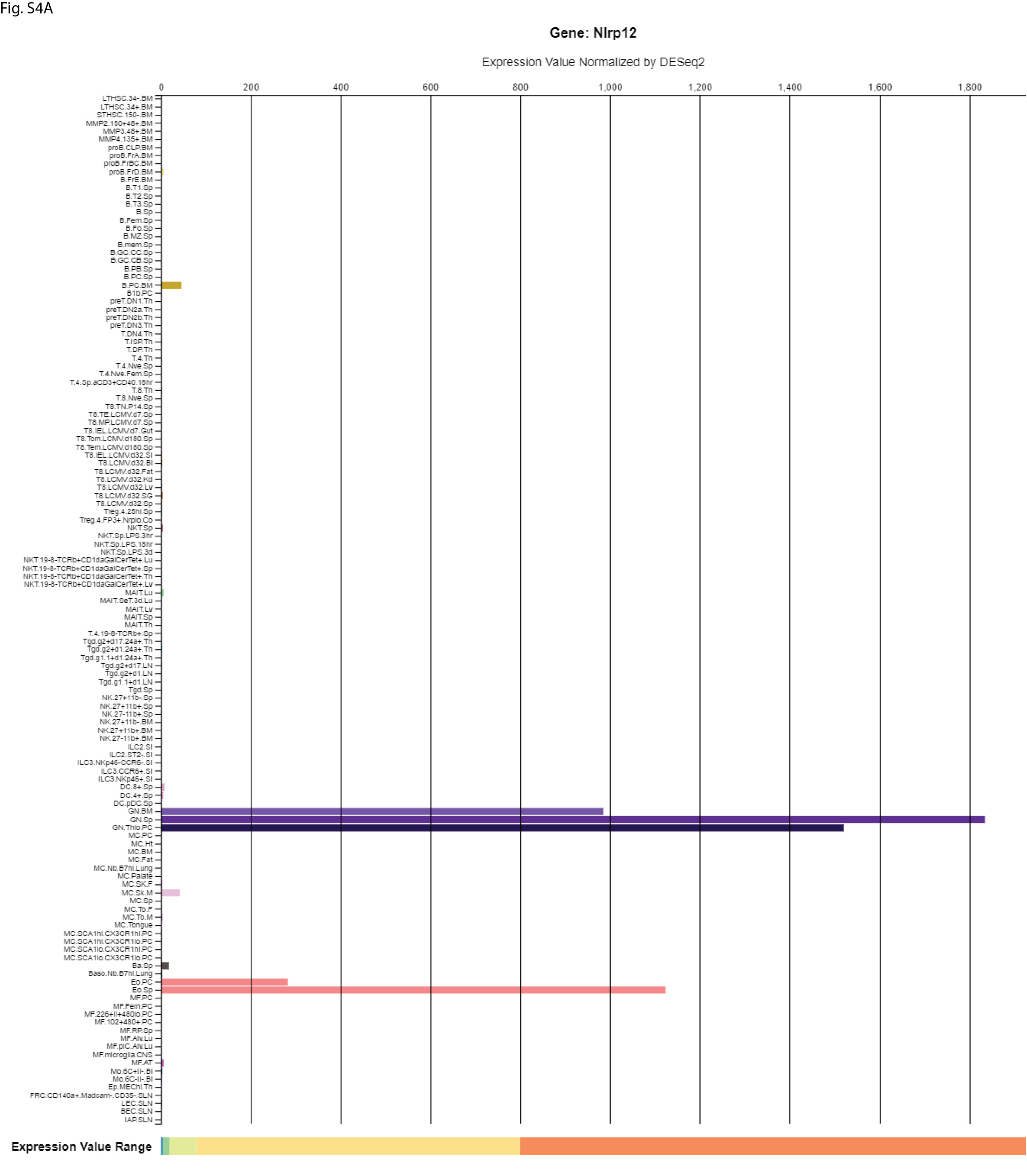

### Supplemental Figure 4B

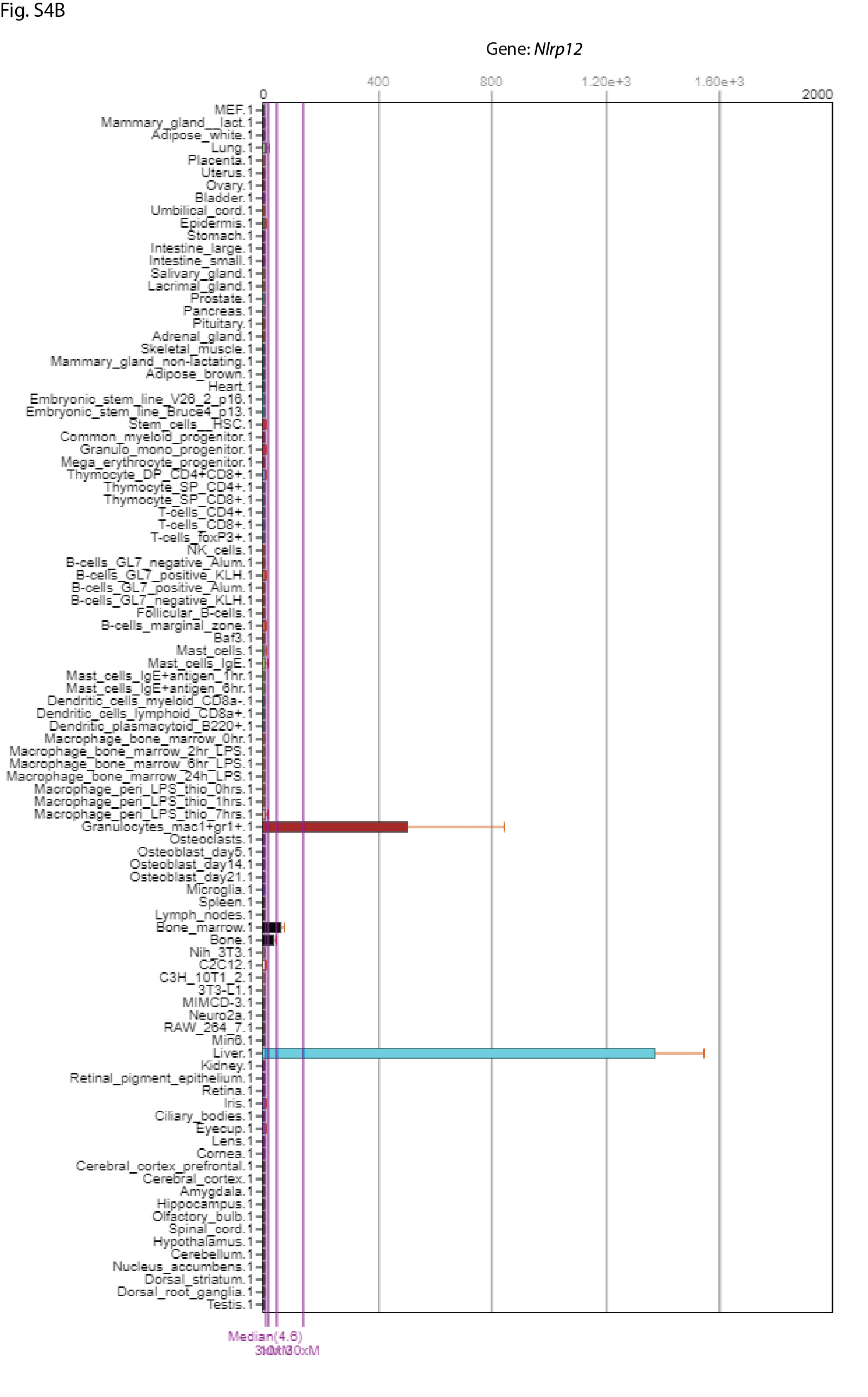

### Supplemental Figure 4C

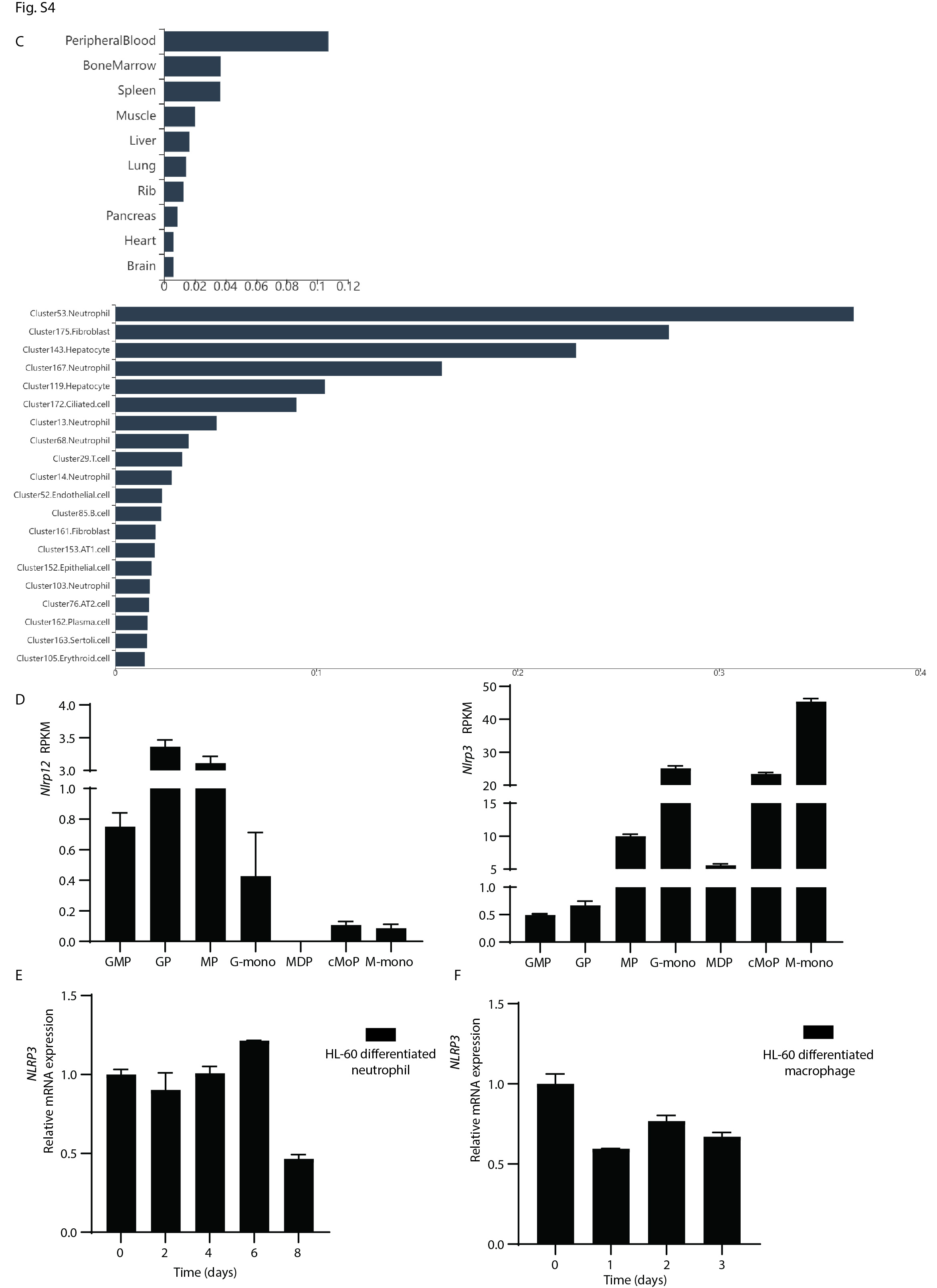

### Supplemental Figure 4G

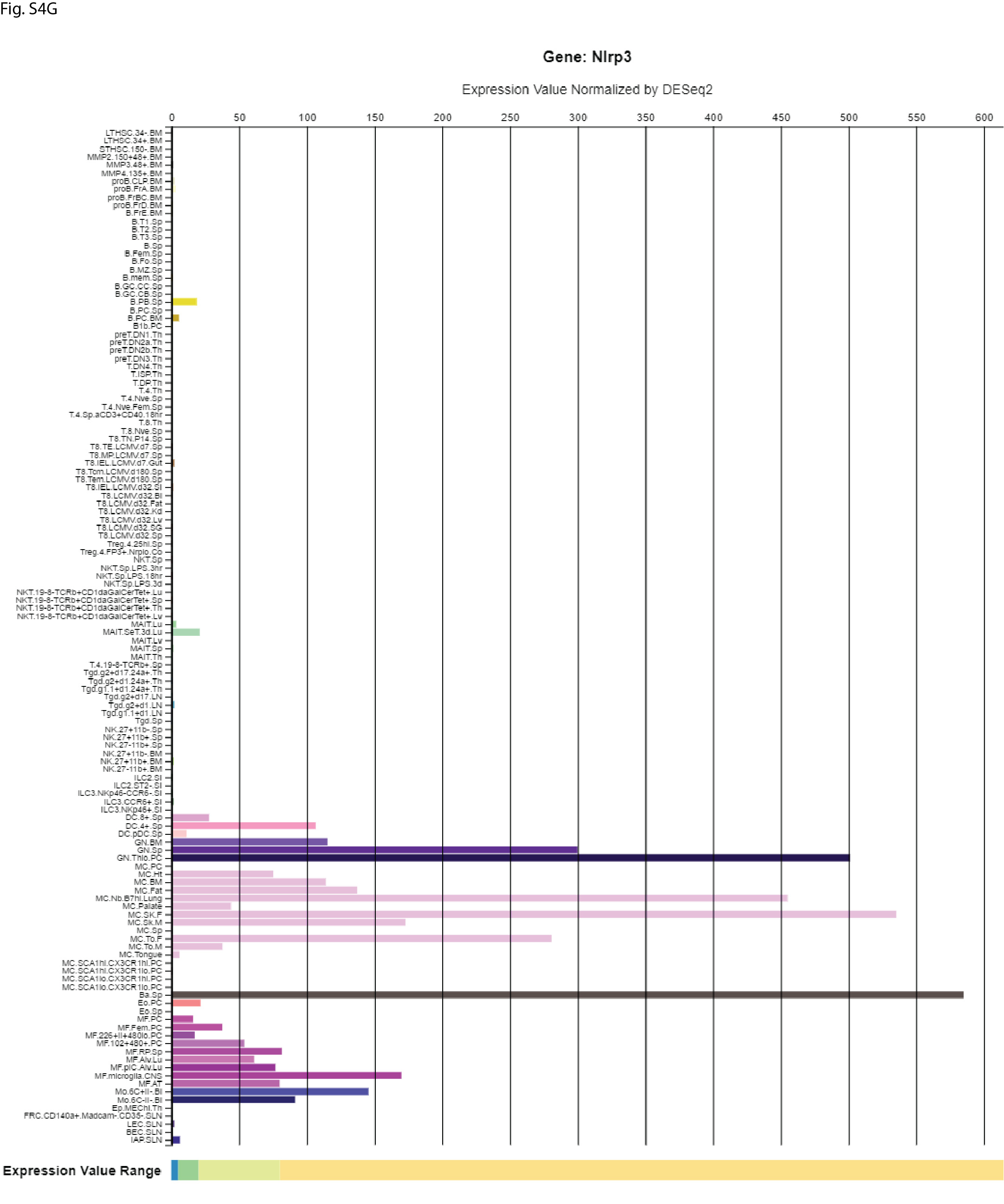

### Supplemental Figure 5

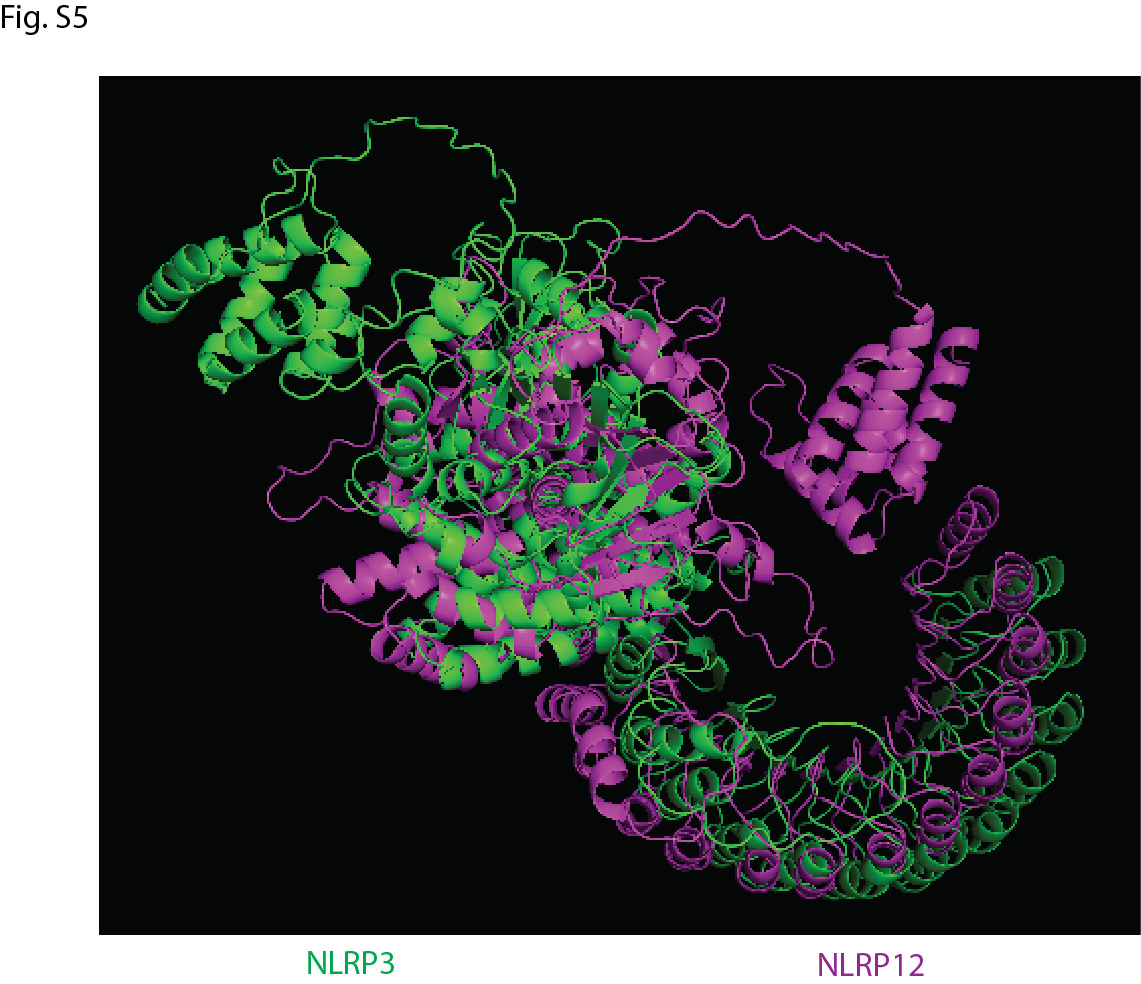

### Supplemental Figure 46

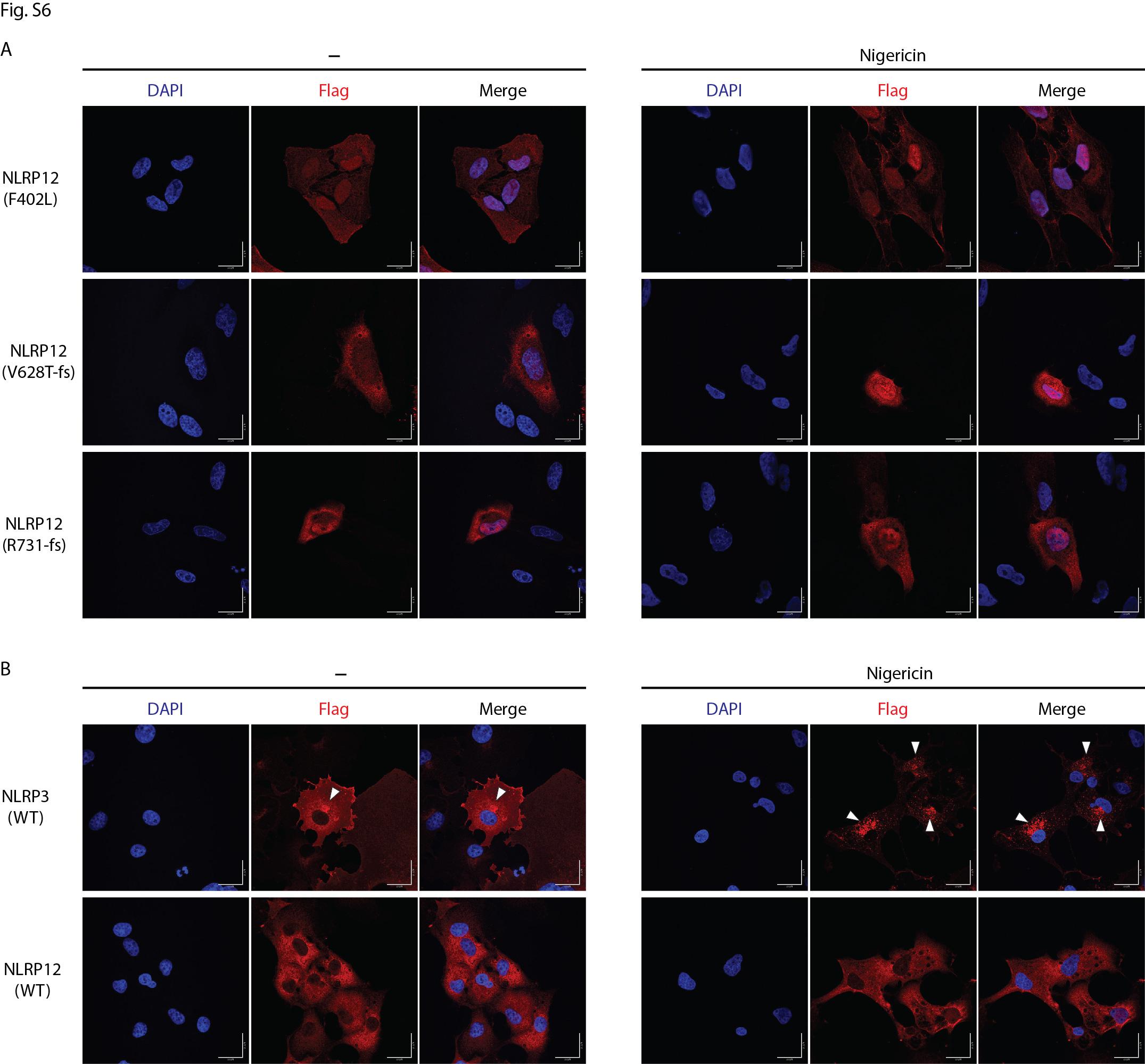

### Supplemental Table 1

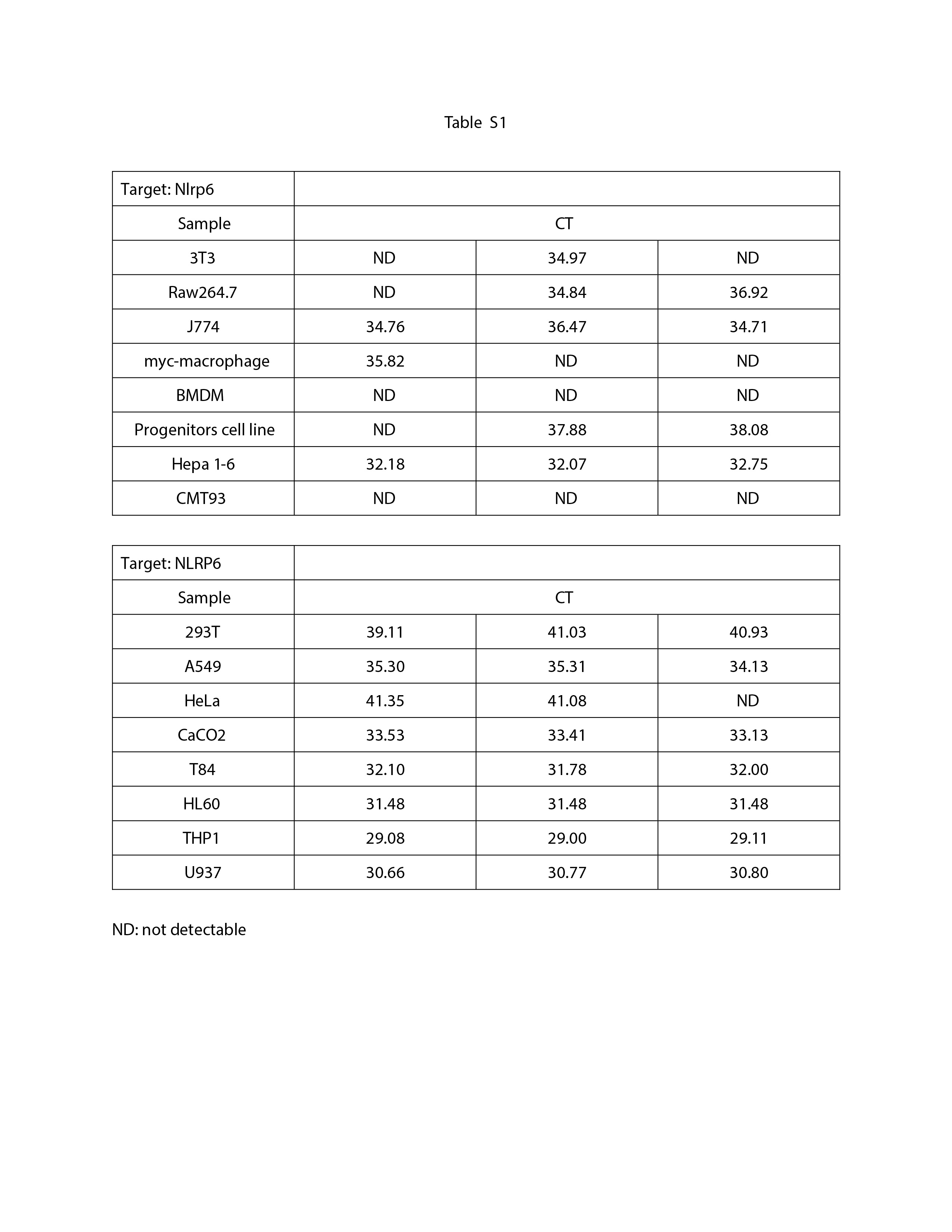
